# A Genomically and Clinically Annotated Patient Derived Xenograft (PDX) Resource for Preclinical Research in Non-Small Cell Lung Cancer

**DOI:** 10.1101/2022.03.06.483171

**Authors:** Xing Yi Woo, Anuj Srivastava, Philip C. Mack, Joel H. Graber, Brian J. Sanderson, Michael W. Lloyd, Mandy Chen, Sergii Domanskyi, Regina Gandour-Edwards, Rebekah A. Tsai, James Keck, Mingshan Cheng, Margaret Bundy, Emily L. Jocoy, Jonathan W. Riess, William Holland, Stephen C. Grubb, James G. Peterson, Grace A. Stafford, Carolyn Paisie, Steven B. Neuhauser, R. Krishna Murthy Karuturi, Joshy George, Allen K. Simons, Margaret Chavaree, Clifford G. Tepper, Neal Goodwin, Susan D. Airhart, Primo N. Lara, Thomas H. Openshaw, Edison T. Liu, David R. Gandara, Carol J. Bult

## Abstract

Patient-derived xenograft models (PDXs) are an effective preclinical *in vivo* platform for testing the efficacy of novel drug and drug combinations for cancer therapeutics. Here we describe a repository of 79 genomically and clinically annotated lung cancer PDXs available from The Jackson Laboratory that have been extensively characterized for histopathological features, mutational profiles, gene expression, and copy number aberrations. Most of the PDXs are models of non-small cell lung cancer (NSCLC), including 37 lung adenocarcinoma (LUAD) and 33 lung squamous cell carcinoma (LUSC) models. Other lung cancer models in the repository include four small cell carcinomas, two large cell neuroendocrine carcinomas, two adenosquamous carcinomas, and one pleomorphic carcinoma. Models with both *de novo* and acquired resistance to targeted therapies with tyrosine kinase inhibitors are available in the collection. The genomic profiles of the LUAD and LUSC PDX models are consistent with those observed in patient tumors of the same tumor type from The Cancer Genome Atlas (TCGA) and to previously characterized gene expression-based molecular subtypes. Clinically relevant mutations identified in the original patient tumors were confirmed in engrafted tumors. Treatment studies performed for a subset of the models recapitulated the responses expected based on the observed genomic profiles.

**Significance:** The collection of lung cancer Patient Derived Xenograft (PDX) models maintained at The Jackson Laboratory retain both the histologic features and treatment-relevant genomic alterations observed in the originating patient tumors and show expected responses to treatment with standard-of-care agents. The models serve as a valuable preclinical platform for translational cancer research. Information and data for the models are freely available from the Mouse Models of Human Cancer database (MMHCdb, http://tumor.informatics.jax.org/mtbwi/pdxSearch.do).

## Introduction

Lung cancer is the leading cause of cancer deaths worldwide (1). Genome-wide analyses have demonstrated that non-small cell lung cancer (NSCLC) differs from most other cancer types quantitatively and qualitatively for its high level of mutational burden and genomic complexity. Further, the two major histologic subtypes of NSCLC: lung adenocarcinoma (LUAD) and lung squamous cell carcinoma (LUSC) (2,3). LUAD and LUSC tumors have distinctive genomic alteration signatures, pathway disruption, and immune host response. Transcriptional subtypes for both LUAD and LUSC have been reported that are associated with differences in patient prognosis, response to treatment, and survival (4,5).

Genomic characterization of tumors has been instrumental in precision medicine strategies for NSCLC through the identification of “druggable” oncogene drivers which, in turn, has expanded treatment options and a growing number of targeted therapy approaches (6). A prominent example of molecularly-guided therapy in NSCLC relates to the finding of activating mutations in the epidermal growth factor receptor (*EGFR*) gene, resulting in constitutive, ligand-independent receptor activity and a high degree of sensitivity to EGFR-targeted tyrosine kinase inhibitors (TKIs) (7). The efficacy of *EGFR* TKIs has been demonstrated in a large number of clinical trials (8). Similar findings have been shown with targeted therapies in patients with an array of other genomically-defined subtypes, such as *ALK-EML4* and *ROS1* fusions, among others (9). Although advances in targeted therapies for NSCLC have transformed treatment options, not all patients respond to treatment and the development of acquired resistance is almost universal. Although resistance mechanisms in some treatment settings for oncogene-driven NSCLC are well established, such as development of the T790M “gatekeeper” mutation after therapy with first and second generation *EGFR* TKIs (10), resistance mechanisms are much more complex in most other therapeutic settings, generally characterized as either secondary mutations or bypass mechanisms. Testing novel treatment strategies and new therapeutic agents to overcome acquired resistance remains a high priority for translational cancer research.

Human tumors engrafted into transplant-compliant recipient mouse hosts (Patient-Derived Xenografts, PDX) retain critical biological properties of a patient’s tumor, including tumor heterogeneity and genomic complexity (11). PDXs have demonstrated utility as preclinical models for testing therapeutic strategies for many cancers, including lung cancer. Previous studies have demonstrated that lung cancer PDX models recapitulate faithfully many aspects of the original patient tumor for histology, karyotype, and genomics (12,13). Lung PDX models have demonstrated the capacity to recapitulate expected sensitivity and resistance patterns to targeted therapies, including clinical responses observed in patients. These models have provided insights into therapies based on other molecular markers (14,15). Collections of PDX models have allowed further studies on understanding the contributing factors affecting engraftment rates, new treatment combinations for lung cancer models which developed resistance, and discovery of new biomarkers for lung cancer treatment (16,17).

In collaboration with the University of California Davis Comprehensive Cancer Center and Northern Light Eastern Maine Medical Center, we generated and characterized (18) a repository of 79 genomically and clinically annotated lung cancer PDX models to use as a platform to support basic research on mechanisms of treatment response and to facilitate translational pre-clinical and co-clinical trial research. This repository is comprised of PDX models that were generated using the NOD.Cg-*Prkdc^scid^ Il2rg^tm1Wjl^*/SzJ (NSG) mouse strain as the host and includes models of high clinical relevance, including *EGFR*- and *KRAS*-mutated lung adenocarcinomas (LUAD) and PI3K-mutant lung squamous cell carcinomas (LUSC). Clinical demographic information, histology, immunohistochemistry images, summarized genomic data, and treatment response data for these PDX models are freely available from The Jackson Laboratory (JAX) PDX web portal hosted by the Mouse Models of Human Cancer database (MMHCdb, http://tumor.informatics.jax.org/mtbwi/pdxSearch.do) (19) and from PDX Finder, a global catalog of thousands of PDX models (20).

## Materials and Methods

### Establishing xenografts

An overview of the PDX model generation process is shown in Fig. 1. All animal procedures were performed at The Jackson Laboratory Sacramento facility under IACUC protocol 12027. NOD.Cg-*Prkdc^scid^ Il2rg^tm1Wjl^*/SzJ (NSG; JAX Stock 005557) animals were housed in individually ventilated polysulfone cages with HEPA filtered air at a density of up to five mice per cage. Cages were changed every two weeks. The animal rooms were on a 12 h light/dark cycle (6 am to 6 pm) with fluorescent lighting. The temperature and relative humidity in the animal rooms were 22 ± 4°C and 50 ± 15%, respectively with 15 air exchanges per hour. Filtered tap water, acidified to a pH of 2.5 to 3.0, and custom LabDiet 5K52 were provided *ad libitum*. Mice were housed in rooms classified as “pathogen and opportunistic-free”. The list of excluded organisms in these rooms and health reports are available on The Jackson Laboratory website (https://www.jax.org/jax-mice-and-services/customer-support/customer-service/animal-health).

**Figure 1.**
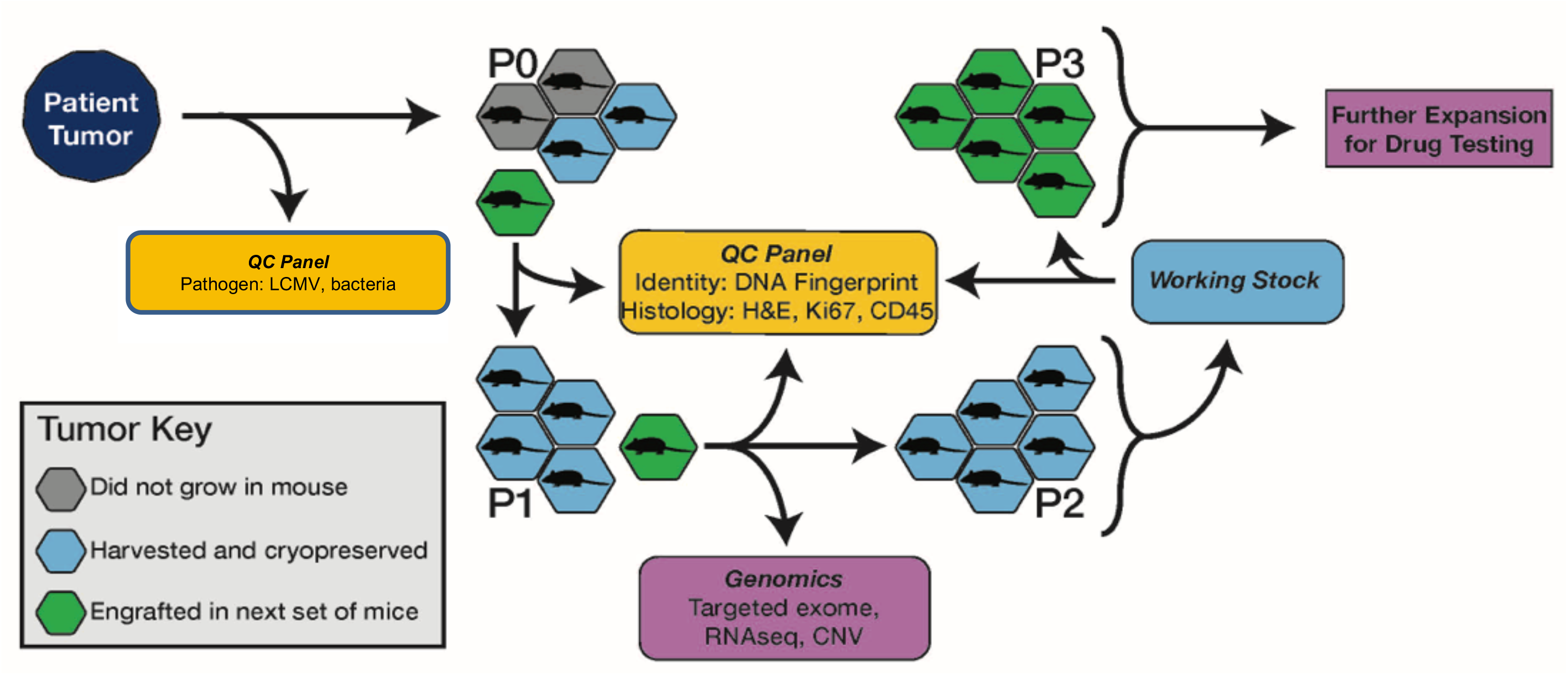
Overview of PDX model generation and characterization for the JAX PDX Resource

Tumor samples from biopsies, pleural effusions, or surgical resections were obtained from NSCLC patients and implanted subcutaneously by trocar in the right flank of up to five, 6-8 week-old female NSG mice without intervening *in vitro* culturing of the tumor cells. Patients were consented by the donating institution to allow unrestricted use of the models and associated data. Most tumors were implanted within 24 hours of surgery and the maximum post-surgery time allowed for implantation was 48 hours. Solid tumors were divided into 3-5 mm^3^ fragments in RPMI medium before implantation. Pleural effusion samples were centrifuged, and the supernatant was removed with a pipet. Pellets were then re-suspended in Dulbecco’s Phosphate Buffered Saline (DPBS), and 200 μl were implanted subcutaneously into the host mouse as a 1:1 bolus of pleural effusion cells in RPMI media and growth factor free Matrigel. Matrigel was not used for subsequent passages.

Once an implanted tumor reached 2000 mm^3^, it was harvested and subdivided into 3-5 mm^3^ fragments which were implanted into five, 6-8 week old female NSG mice for expansion to P1. For quality control assessment (see below), a 50 mm^3^ fragment was collected in 10% neutral buffered formalin and a formalin-fixed, paraffin-embedded block was generated. The remaining fragments were cryopreserved in 10% DMSO. When P1 tumors reached ~2000mm^3^, they were harvested and subdivided into 3-5 mm^3^ fragments which were subsequently embedded in FFPE for quality control, snap-frozen for genomics, placed into RNALater (Ambion) for RNA-Seq, and viably cryopreserved in 10% DMSO.

To establish cohorts of tumor-bearing mice for drug treatment studies, 3-5 mm^3^ tumor fragments or 40 μl of minced tumor were subcutaneously implanted in the right rear flank of NSG mice by trocar or a 14-gauge disposable needle. Low passage tumor fragments (P3-P6) were used to establish cohorts of tumor-bearing animals for dosing studies. Tumor volumes were monitored with ULTRA-Cal IV digital calipers (Fowler, Newton, MA). Individual tumor-bearing mice were randomized into treatment cohorts of 8-12 mice each on an accrual (asynchronous growth) basis once individual tumors reached an initial volume of approximately 100-300 mm^3^. For some studies, tumors were removed and divided, with half of the material preserved in neutral-buffered formalin and half flash-frozen.

### PDX model quality control

The quality control procedures employed for PDX models included testing the patient tumor for LCMV (lymphocytic choriomeningitis virus) and bacterial contamination. The engrafted tumors at P0 and P1 were DNA fingerprinted using a Short Tandem Repeat (STR) assay (21) and then compared to the profile of the patient sample to ensure correct tissue provenance. Immunohistochemistry (IHC) for human CD45 antibodies (IR75161-2, Agilent Technologies) was performed on FFPE blocks of engrafted tumors to identify cases of lymphomagenesis, which have been reported previously in PDXs (22). IHC for human Ki67 (IR62661-2, Agilent Technologies) was used to ensure the propagated tumors were comprised of human cells. H&E sections of engrafted tumors were evaluated by a board-certified pathologist (RGE) to verify the concordance of the morphological features of the engrafted tumor to the patient tumor. Engrafted tumors were assessed by sequencing or digital droplet PCR to ensure that diagnostic/therapeutic molecular markers identified in patient samples were present in the engrafted tumors (Supplementary Table S1).

### Genomic characterization of engrafted tumors

The genomes of engrafted tumors were characterized at either the P0 or P1 passage (Fig. 1). Sequencing using The Jackson Laboratory (JAX) Cancer Treatment Profile (CTP) targeted gene panel (23) was performed to determine point mutations, single nucleotide polymorphisms insertions/deletions (indels). The JAX CTP panel comprises 358 genes, 190 of which are clinically actionable genes for cancer treatment. Transcriptome analysis was conducted using expression microarrays or RNA sequencing (RNA-Seq). RNA-Seq data was also used to identify putative translocations, RNA splicing abnormalities, and fusion transcript events. Sequencing data were analyzed using Xenome to separate mouse sequences from human-derived sequence reads (24). Copy Number Variants (CNVs) were determined using the Affymetrix Human SNP 6.0 array. Detailed protocols for nucleic acid extraction, library preparation, and data analysis are described elsewhere (18). The JAX Clinical Knowledge Base (CKB) database was used to annotate variants and gene expression with clinical relevance (25). Clinical information and analyzed genomic data of the PDX models were then uploaded to a public PDX web portal hosted by the Mouse Models of Human Cancer database (MMHCdb).

### Analysis of genomic data from engrafted tumors

Mutation and copy number (CN) analysis and quantification of gene expression for all engrafted tumor samples were performed as described in Woo *et al*. (18). Tumor mutation burden (TMB) and microsatellite instability (MSI) were estimated for each tumor sample that was characterized using the JAX CTP targeted gene panel. TMB was calculated using variants that met all quality criteria (coverage, strand bias, mapping quality, and read rank position) and were not present on a curated list of false-positive variants (loci prone to sequencing and analysis errors and/or associated with highly polymorphic genes: *MUC4*, *MUC5B*, *MUC16*, *MUC17*, and *HLA-A*). Only likely somatic mutations based on germline filtering criteria that were predicted with high or moderate functional impact (i.e., non-synonymous changes, frame shifts, stop losses/gains, and splice-site acceptor/donor changes) were retained. TMB was estimated by dividing the number of variants that met the quality criteria by the length (in Mb) of the CTP gene panel. High TMB was defined as 22 mutations/Mb, which was calculated based on the TMB distribution of all PDX models analyzed as follows: Q3 (third quartile of TMB) + 1.5 x inter-quartile range of TMB. The MSIsensor2 (26) algorithm (https://github.com/niu-lab/msisensor2) was used to determine the MSI status of JAX samples. The samples with MSI-Percentage > 20% were considered MSI-High. This threshold demonstrates good differentiation between MSI-High (MSI-H) and MSI-Stable (MSI-S) samples during MSIsensor2 algorithm development and internal benchmarking.

To summarize the mutations prevalent in the PDX models, oncoplots for LUAD and LUSC were created using all the somatic and clinically relevant point mutations and indels in the models. All mutations were concatenated across the PDX samples for models with more than one sample sequenced and derived a unique list of mutations per model. The list of mutations was converted to annovar format, and the table_annovar.pl utility from annovar (Version date: 2014-11-12) with the parameters (-buildver hg38 -remove -protocol refGene,cytoBand,exac03,avsnp147,dbnsfp30a -operation gx,r,f,f,f -nastring. -polish - otherinfo) (27) was used to convert the variants to Mutation Annotation Format (MAF). Finally, the oncoplot function from maftools v2.2.10 (28) was used to create the oncoplot for genes which were mutated >30% frequency of the models. Gene mutation frequencies of LUAD and LUSC in the TCGA PanCancer Atlas were obtained from cBioPortal (29) for comparison with the PDX data. Genes were classified as oncogenes or tumor suppressor genes (TSGs) based on OncoKB annotations (Download date: 2020/9/17) (30).

Copy number data were visualized using GenVisR (31) on a per-sample basis and an overall gain and loss frequency basis within the LUAD and LUSC groups. For frequency calculation, one sample was selected to represent each model, and log_2_(CN ratio) = ±0.5 was used as a cut-off to call CN gain and loss.

To summarize the expression data, the percentile rank z-score values from stranded RNA-Seq, non-stranded RNA-Seq, and microarray platforms were combined and a correlation heatmap was plotted using the Pretty Heatmaps package in R (https://cran.r-project.org/web/packages/pheatmap/index.html).

### Gene expression-based subtypes of LUAD and LUSC PDXs

To determine if previously identified molecular subtypes for LUAD (TRU, PIF, and PPR) and LUSC (CLA, PRI, BAS, and SEC) were represented in our repository of lung cancer PDX models, we used Nearest Template Prediction (NTP) implemented in the R package CMScaller (32). We selected genes from publicly available RNA-seq data to enrich for classification of the subtypes, following similar methods as those recently used to develop a classifier for colorectal cancer. TCGA raw gene expression data (non-stranded RNA-Seq) were downloaded from the Broad GDAC Firehose repository for LUAD (7) and LUSC (33). Molecular subtype annotations for LUAD and LUSC samples from TCGA were downloaded using the TCGAquery_subtype function in the R package TCGAbiolinks (34). Gene expression data were harmonized between TCGA and PDX samples and filtered for lowly expressed genes (mean normalized expression ≤ 1), so that only genes expressed in PDX samples were used for training the classifier. We estimated differential expression between subtypes in TCGA samples using the R package DESeq2 v. 1.28.1 (35). Genes were classified as differentially expressed for each subtype using a threshold of an FDR-adjusted P-value of ≤ 0.01 and an absolute log_2_ fold-change value > 1. The lists of genes identified by these criteria as differentially expressed were compared. All differential genes that showed discrimination between at least one subtype and the others (e.g., not commonly identified as differentially expressed among all subtypes) were included to generate custom templates for NTP. We also selected genes for template creation that showed a large range of expression values among LUAD and LUSC cell lines in the Cancer Cell Line Encyclopedia (36) and highly expressed in at least a subset of the cell lines. Finally, we selected genes to exclude from the training set by performing a differential expression analysis between matched lung cancer patients (7 samples) and PDX samples (41 samples) obtained from the NCI Patient-Derived Model Repository (PDMR, NCI-Frederick, Frederick National Laboratory for Cancer Research, https://pdmr.cancer.gov/), using DESeq2. The genes with an absolute log_2_ fold change ≥ 1 were excluded from the model training set as they may be lost upon engraftment.

The final list of genes used to train our NTP classifiers included 3,525 genes for LUAD and 3,544 genes for LUSC. Raw expression data from TCGA for these genes were used to generate templates for NTP. TCGA samples were split randomly into 80% training and 20% validation sets stratified by labeled subtype. Custom templates were then prepared for LUAD and LUSC training sets using the functions subDEG and ntpMakeTemplates in the R package CMScaller. Subtype prediction performance was estimated using the validation sets with the function ntp in the R package CMScaller, specifying 1000 permutations. Performance of subtype predictions from NTP was calculated using 20% of the labeled TCGA data held back for validation. For each subtype, the performance of the NTP classification was measured by precision, recall and F1-score. The overall accuracy of the predictions was calculated as the unweighted average of the proportion of true positives to samples of each subtype. Finally, we used the custom templates to generate subtype predictions for the unlabeled PDX models with non-stranded RNA-Seq data, and we considered high confidence subtype classifications with FDR-adjusted P-values < 0.05.

### PDX Treatment Studies

Tumor-bearing mice at low passage (P3-P6) were assigned to cohorts (8-10 mice per treatment group) and treated with single and combination agent therapies depending on the lung cancer subtype and presence of targetable molecular markers. Vehicle treated mice were used as controls. Treatment was initiated when tumors reached approximately 70-300 mm^3^. Tumors were monitored until the end of the dosing study (typically 28 days) or when the tumors reached 2000 mm^3^. To monitor for toxicity effects from treatment, animal body weight was monitored three times weekly throughout the study, and percent body weight loss was calculated for each mouse. Animals with >20% body weight loss were euthanized and recorded as treatment-related deaths. Tumor volumes were calculated from digital caliper raw data by using the formula:

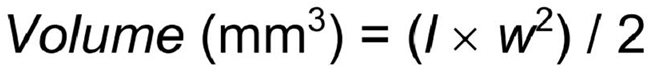

The value *w* (width) was assigned as the smaller of two perpendicular tumor axes and the value *l* (length) as the larger.

The response to treatment was classified using modified Response Evaluation Criteria In Solid Tumors (RECIST) criteria and by Tumor Growth Inhibition (TGI). The RECIST criteria were established using the percentage of tumor volume change (ΔVol) at the final study day (i.e., seven days after the last treatment) compared with the baseline tumor volume at Day 0 or Day 1. A response classification was adapted from Gao *et al*. (37) as follows:

Step 1: Calculate percent change in tumor volume for each animal as V: ((end_volume – start_volume)/start_volume) * 100) at day 21
Step 2: Within each group, find the minimum V as Vm
Step 3: Within each group, find the mean (average) V as Va
Step 4: Determine modified RECIST category as shown:

Complete response (CR): Vm < −95%, Va < −40%
Partial response (PR): Vm < −50%, Va < −20%
Stable disease (SD): Vm < 35%, Va < 30%
Progressive disease (PD): Anything else

Tumor growth inhibition (TGI) was summarized as the % of tumor volume change in treatment arms relative to the control. The % TGI is defined as (1 – (mean volume of treated tumors)/(mean volume of control tumors)) × 100% at study termination. Graphical summaries of treatment responses for each cohort were generated with custom visualization software (https://github.com/TheJacksonLaboratory/PDX-SOC) and are available from the PDX data portal on MMHCdb.

### Western blots

Immunoblotting was performed on treated tumors using methods described previously (38).

## Results

### Enrollment and patient characteristics

The clinical and demographic data for the patients from whom the lung PDX models were generated are summarized in Table 1. The median age for patients was 63 (range 42-85). Slightly more female (n=44) versus male patients (n=35) are represented in the patient population from which the models were derived. Most of the patients (80%) reported their race as White. The self-reported smoking status of the patient cohort was as follows: former (44%), current (20%), and never (8%). Most of the patients (30 of 33) diagnosed with LUSC were treatment naïve at the time their tumor tissue was acquired to generate a PDX model. For LUAD, half of the patients (19 of 37) were treatment-naïve at the time of PDX model generation.

**Table 1.**
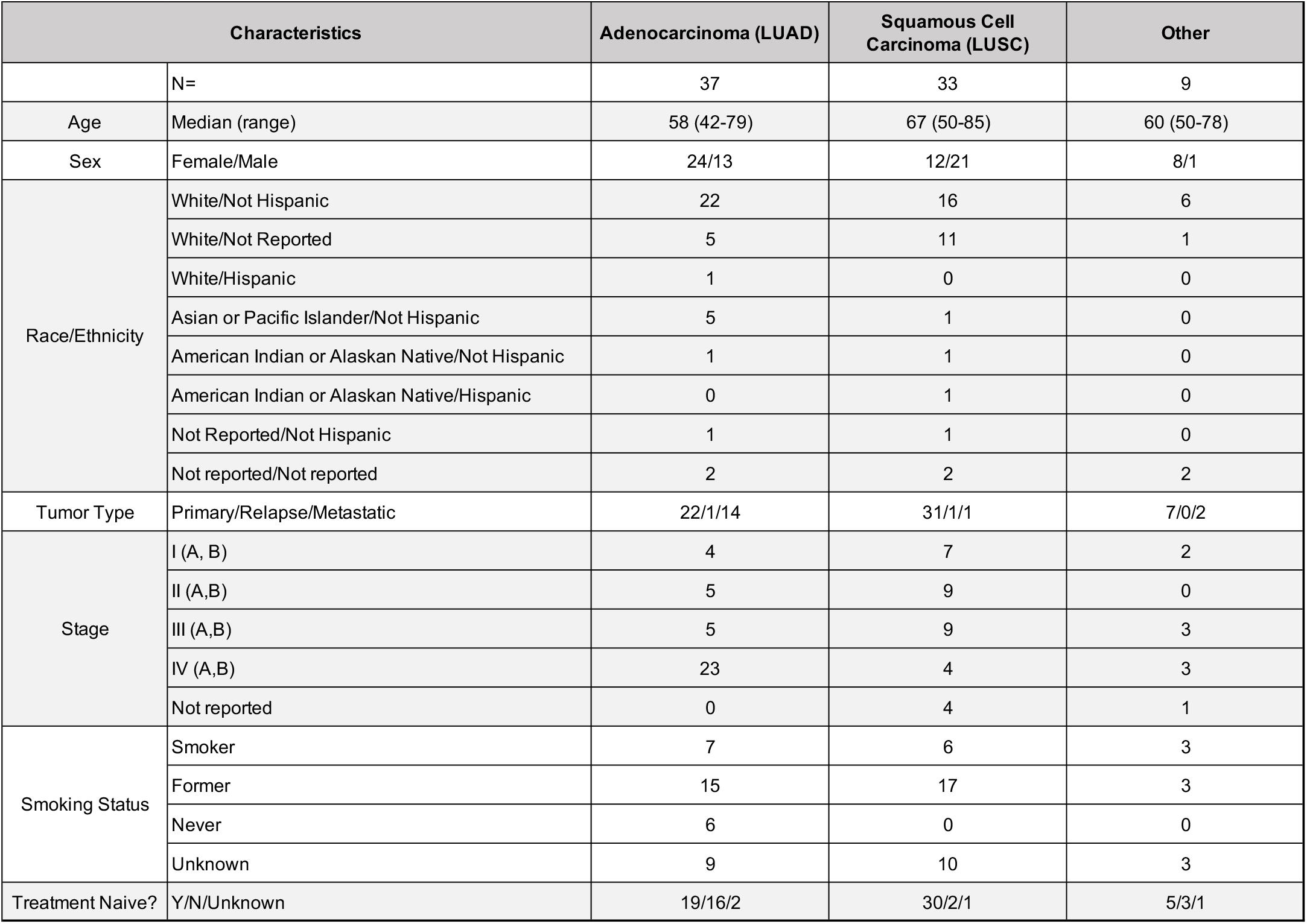
Summary of clinical and demographic data for patients whose tumor material was used to generate the JAX collection of lung cancer PDX models.

| Characteristics |  | Adenocarcinoma (LUAD) | Squamous Cell Carcinoma (LUSC) | Other |
| --- | --- | --- | --- | --- |
|  | N= | 37 | 33 | 9 |
| Age | Median (range) | 58 (42-79) | 67 (50-85) | 60 (50-78) |
| Sex | Female/Male | 24/13 | 12/21 | 8/1 |
| Race/Ethnicity | White/Not Hispanic | 22 | 16 | 6 |
|  | White/Not Reported | 5 | 11 | 1 |
|  | White/Hispanic | 1 | 0 | 0 |
|  | Asian or Pacific Islander/Not Hispanic | 5 | 1 | 0 |
|  | American Indian or Alaskan Native/Not Hispanic | 1 | 1 | 0 |
|  | American Indian or Alaskan Native/Hispanic | 0 | 1 | 0 |
|  | Not Reported/Not Hispanic | 1 | 1 | 0 |
|  | Not reported/Not reported | 2 | 2 | 2 |
| Tumor Type | Primary/Relapse/Metastatic | 22/1/14 | 31/1/1 | 7/0/2 |
| Stage | I (A, B) | 4 | 7 | 2 |
|  | II (A,B) | 5 | 9 | 0 |
|  | III (A,B) | 5 | 9 | 3 |
|  | IV (A,B) | 23 | 4 | 3 |
|  | Not reported | 0 | 4 | 1 |
| Smoking Status | Smoker | 7 | 6 | 3 |
|  | Former | 15 | 17 | 3 |
|  | Never | 6 | 0 | 0 |
|  | Unknown | 9 | 10 | 3 |
| Treatment Naive? | Y/N/Unknown | 19/16/2 | 30/2/1 | 5/3/1 |

### PDX Models

A summary of the 79 lung PDX models described in this report is presented in Supplementary Table S1. The collection of models is comprised mostly of lung adenocarcinomas (37 models) and lung squamous cell carcinomas (33 models). Models for other lung cancer types available from the repository include four small cell carcinomas, two large cell neuroendocrine carcinomas, two adenosquamous carcinomas, and one pleomorphic carcinoma. Only the LUAD and LUSC models were used in the analyses described in this report. On average, 38% of implantations resulted in successful engraftment, similar to other reports on lung cancer PDXs (34 – 39%) (16). Information about the PDX models, including clinical information, genomic data, and treatment response studies, are available from the Mouse Models of Human Cancer database (MMHCdb).

### Quality Control (QC)

All patient tumor samples were negative for LCMV. All engrafted tumors demonstrated positive labeling for human Ki67 protein. Results of STR analysis for each model confirmed the engrafted tumor originated from the expected patient tumor. All models for which hematoxylin and eosin (H&E) stained slides were available for both patient and engrafted tumors data were determined to have moderate to high concordance following visual evaluation of the images by a board-certified pathologist (RGE). Representative histology images and the pathologist’s notations for the PDX models are available from MMHCdb.

Of 95 tumors engrafted, 13 (16%) were identified as lymphoid tumors based on positive staining for human CD45 antigen. These tumors likely arose from transplanted Epstein-Barr Virus (EBV)-infected human B-cells (39). The corresponding PDX models were removed from the JAX repository resulting in the final set of 79 models described here. A similar percentage of lymphomagenesis was reported in another PDX lung cancer model collection (40).

The Xenome (24) algorithm was used to determine human and mouse origins of sequence data generated from engrafted tumor samples. The average percentage of human sequences was 87% (53% - 99%) and 79% (50% - 89%) for the CTP assay and RNA-Seq, respectively (Supplementary Fig. S1). The average percentage of mouse sequences identified by Xenome for both CTP and RNA-Seq data was ~12% (0.5% - 47%). The percentage of sequence reads that were classified as “both” or “ambiguous” were <0.2% and ~0.6% for both platforms, respectively, and the “neither human nor mouse” category averaged 0.04% for CTP and 7.8% for RNA-Seq. The observed differences in the percentages of mouse and human sequences are likely due to platform differences (sequence capture method for CTP compared to direct sequencing for RNA-Seq). The difference was significant for human sequence reads (Welch Two Sample t-test; p-value = 0.00000002191) but not for sequence reads classified as mouse.

### Genomic characterization: Somatic mutation

Genes on the JAX CTP panel that were mutated in at least 30% of the LUAD and LUSC PDX models are summarized in Fig. 2A and B. The complete gene list with gene mutation frequencies in PDX models of LUAD, LUSC, all lung cancer, and all other cancer types is available in Supplementary Table S2. As has been observed previously in many human cancers, *TP53* is the most commonly mutated gene in both the LUAD and LUSC subtypes of non-small cell lung cancer (7,33). Large genes such as titin (*TTN*), usherin (*USH2A*), and mucins (*MUC4, MUC5B, MUC16, MUC17*) also have high mutation frequencies. As reported previously, it is likely that the high mutation frequencies in these genes are a consequence of their size and do not represent driver mutations (3).

**Figure 2.**
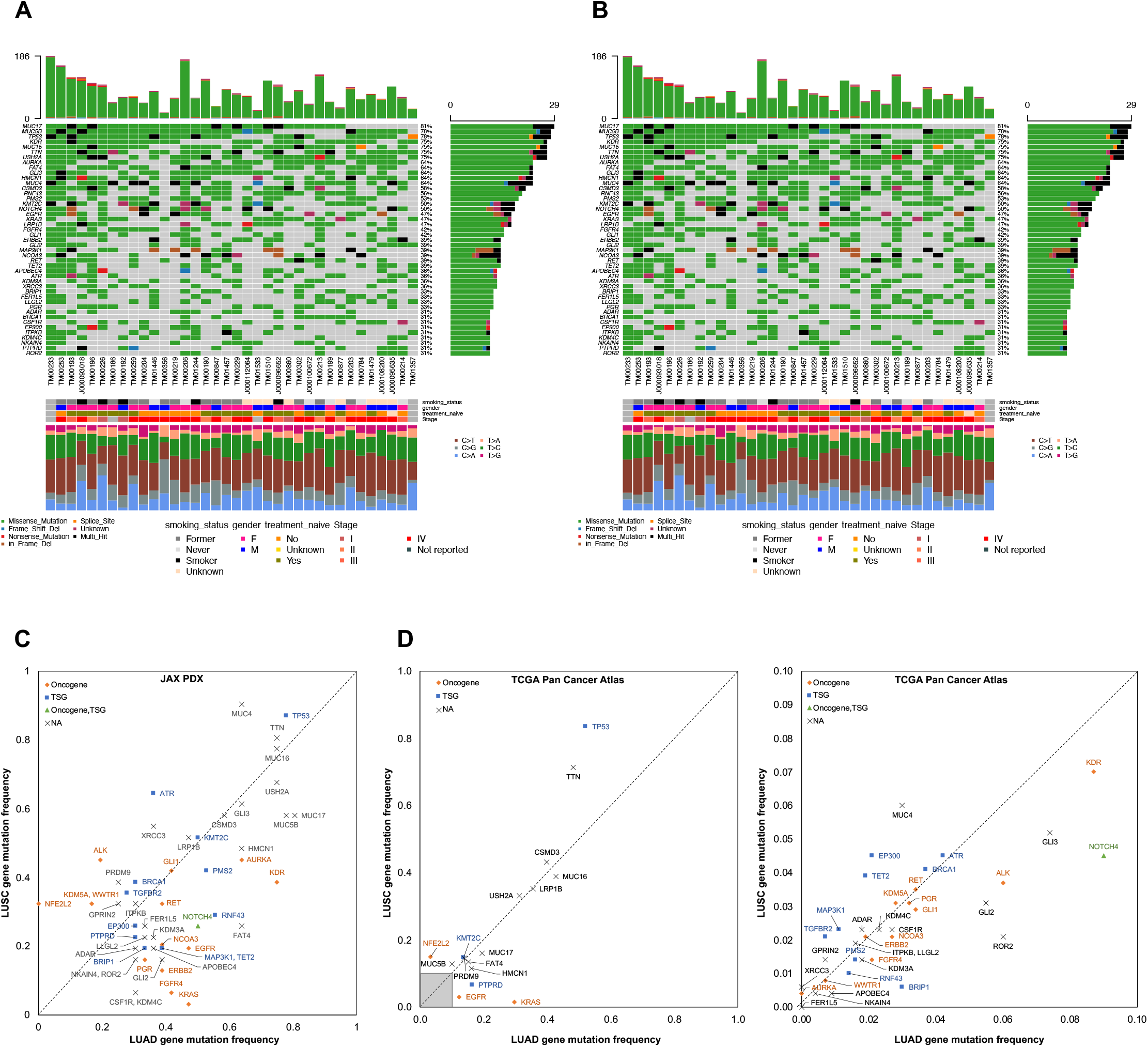
Somatic mutations in lung cancer PDX models **A,B**. Oncoplot of the most frequently mutated genes in LUAD (**A**) and LUSC (**B**) PDX models. The oncoplot shows the PDX models in a horizontal orientation, annotated with smoking status, gender, treatment status and stage of cancer. Genes with mutation frequency > 30% is shown on the vertical axis. The bar plot at the top has the frequency of mutations for each PDX model, while the right bar plot has the frequency of mutations in each gene. Colors in the oncoplot columns indicate different mutation types (see legend for details). The bottom panel shows the classification of the SNPs into transitions and transversions. (**C)** Comparison of gene mutation frequency in LUAD and LUSC PDX models (frequency >30%). (**D)** Comparison of gene mutation frequency in LUAD and LUSC TCGA samples (left: frequency >10%, right: frequency < 10%). Oncogene and Tumor Suppressor Gene (TSG) annotations from OncoKB.

An evaluation of mutation frequencies between adenocarcinomas and squamous cell tumors in the JAX PDX repository revealed that mutation frequencies for some genes are characteristic of the NSCLC subtype (Fig. 2C). Because of the relatively small number of PDX models in this analysis, these trends cannot be considered definitive. However, several of the patterns in the JAX collection are also observed in the TCGA PanCancer Atlas from cBioPortal (29) (Fig. 2D). Genes that are more frequently mutated in LUSC in both the JAX lung PDX and TCGA datasets include *NFE2L2*, *TP53*, and *MUC4*. Genes that show higher mutation frequencies in LUAD in both datasets include *KRAS*, *EGFR*, *NOTCH4*, *HMCN1*, and *MUC17*. Other genes identified as being characteristic of LUAD and LUSC in the two collections that did not overlap included *AURKA*, *FER1L5*, *TET2* and *ALK*. Several factors could explain these differences. First, the JAX PDX samples were sequenced at very high coverage (mean coverage = 941x) compared to the whole-exome sequencing of TCGA samples (~100x) (41). Second, the types of samples used to generate the two resources differ. The tumor types used to generate the JAX resource were often selected by collaborating clinical oncologists based on known clinical (e.g., stage, prior treatment, metastasis, relapse) and/or genomic features. Also, a greater proportion of late-stage tumors (Stage II or later) are found in the JAX PDX models (LUAD: 89%, LUSC: 63%) compared to TCGA PanCancer Atlas (LUAD: 41%, LUSC: 51%) (Supplementary Fig. S2).

Mutations in *EGFR, KRAS*, *ALK*, and *ERBB2* in lung cancer are known to confer sensitivity or resistance to tyrosine kinase inhibitors (TKIs) and other targeted therapies. For 22 of the LUAD PDX models, genomic testing of the patient tumor was provided. All the engrafted tumors retained the clinically relevant mutations of the donor patient’s tumor (Supplementary Table S1). The confirmed mutations include *EGFR* L858R and T790M mutations, *EGFR* exon 19 deletion, *EGFR* exon 20 insertion, *ERBB2* exon 20 insertion, *EML4*-*ALK* fusion, and KRAS G12. For model TM01244, the expected *EGFR* T790M mutation was not observed in the sequence data from the CTP targeted gene panel, but the presence of the mutation was confirmed by droplet digital PCR (ddPCR). The failure of the targeted gene sequencing to identify the mutation in this case could be due to the random sampling of a heterogeneous patient tumor carrying subclonal mutations during the establishment of the PDX model (11,42).

Although patients with LUSC have limited targeted treatment options compared to those diagnosed with LUAD, recent findings of recurrent genomic alterations that are characteristic of this histologic subtype, including activating alterations in *PIK3CA*, *KRAS*, and *MET* that may have therapeutic implications, and provide future therapeutic avenues for research after chemotherapy/immunotherapy options have been exhausted (43). Within the JAX PDX collection, clinically relevant mutations of these genes were found in 25 of the LUSC models (Supplementary Table S1). Only one patient tumor was tested before establishing the PDX model. The KRAS G12C mutation detected in the patient tumor was also observed in the engrafted tumor (TM00231).

Considering both LUAD and LUSC models, 84% (n=69) of the engrafted tumors harbored clinically relevant mutations where clinical relevance was based on annotations from the JAX CKB database (25). We also computed the TMB and MSI for all PDX samples with CTP sequencing data (Supplementary Fig. S3, Supplementary Table S1) as TMB and MSI are used as biomarkers for immunotherapy response (44–48). We observed trends similar to other lung cancer data sets, where lung tumors are rarely MSI-high, but do have high TMB scores (44,48). Within the JAX PDX collection, none of the lung cancer PDX models are MSI-high (MSI score > 20); while ten of the models are classified as high tumor mutation burden (TMB score > 22). The MSI and TMB scores are similar across multiple samples and passages of the same model, indicating that these genomic features are maintained throughout passaging and expansion.

### Genomic characterization: Copy Number Alteration

Recurrent gains and losses of chromosomal regions have been documented in NSCLC previously, including gains in *MYC*, *EGFR*, *CCND1* and losses in *LRP1B* and *CDKN2A* (7,33). In LUSC, gains in 3q and losses in 3p and 5q occur more frequently than in LUAD (49). An overview of the gains and losses observed in engrafted tumors from the JAX NSCLC PDX model collection is provided in Fig. 3; gain and loss frequencies of individual genes are provided in Supplementary Table S3. The frequently amplified and deleted chromosomal regions are consistent with previous studies.

**Figure 3.**
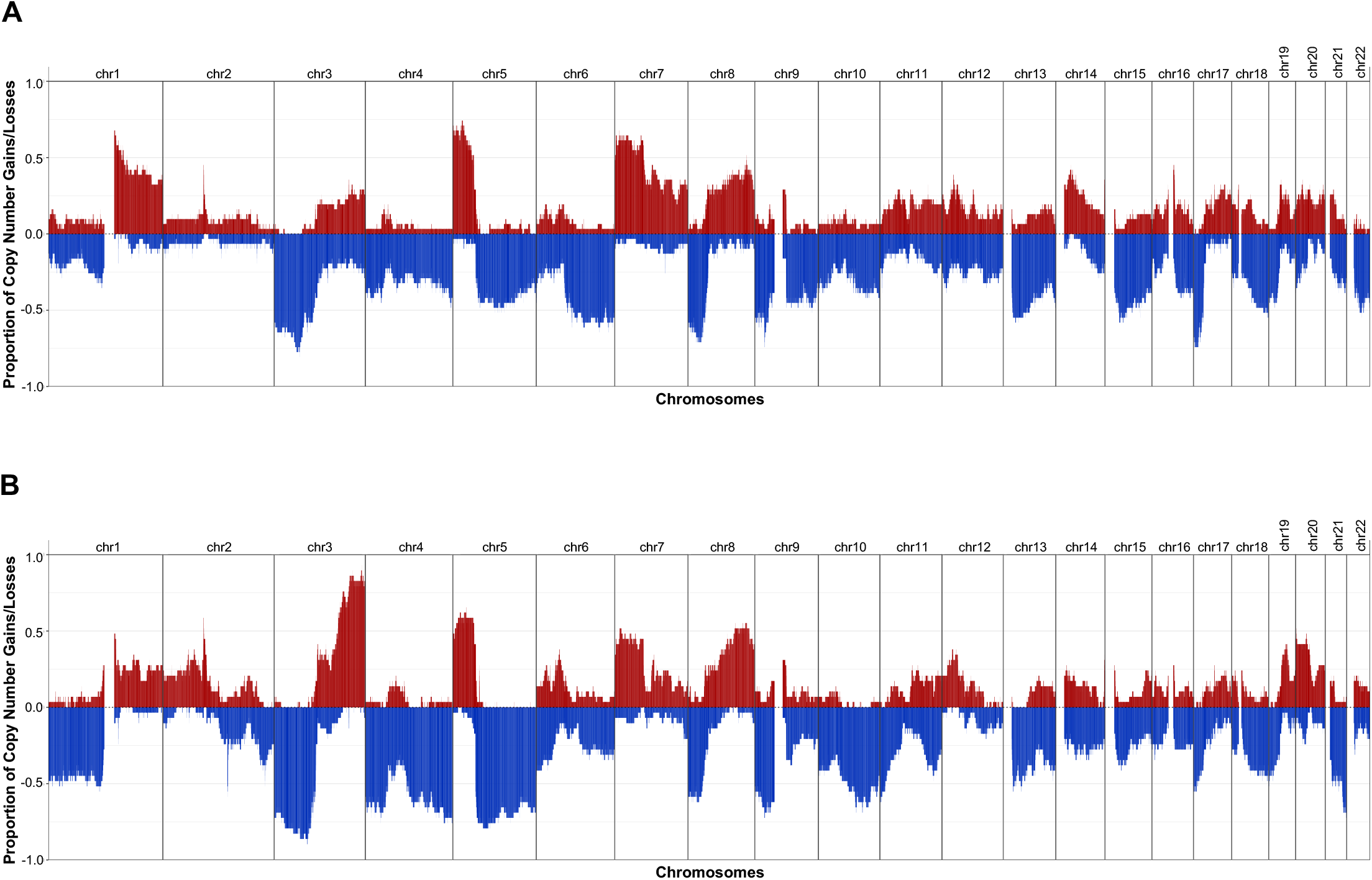
Copy number alterations in lung cancer PDX models **A,B**. Frequency of copy number gain and loss for **(A)** LUAD and **(B)** LUSC PDX models. CN Gain: log2(CN/ploidy) > 0.5; CN Loss: log2(CN/ploidy) < −0.5. One sample per model was used to calculate the frequency.

Copy number profiles of individual PDX samples are shown in Supplementary Fig. S4. Amplifications reported for patient tumors were also observed in the corresponding PDX model. For example, *MET*, *EGFR*, and *MYC* amplifications reported in a patient tumor were recapitulated in the corresponding PDX model (TM00784). Different engrafted tumor samples derived from the same PDX model had high concordance in copy number (11).

### Genomic characterization: Transcriptional profiling

Unsupervised hierarchical clustering of the lung models based on gene expression is shown in Fig. 4A. The samples are clustered primarily by the platform (RNA-Seq or microarray) and then by the diagnosis within each method. The highest similarity is observed between different samples of the same model assayed by the same platform. Tumor samples derived from the same PDX model display a higher correlation in expression, regardless of platforms, than the background of correlating different models (Supplementary Fig. S5), indicating that the expression profile is retained during engraftment, expansion, and passaging.

**Figure 4.**
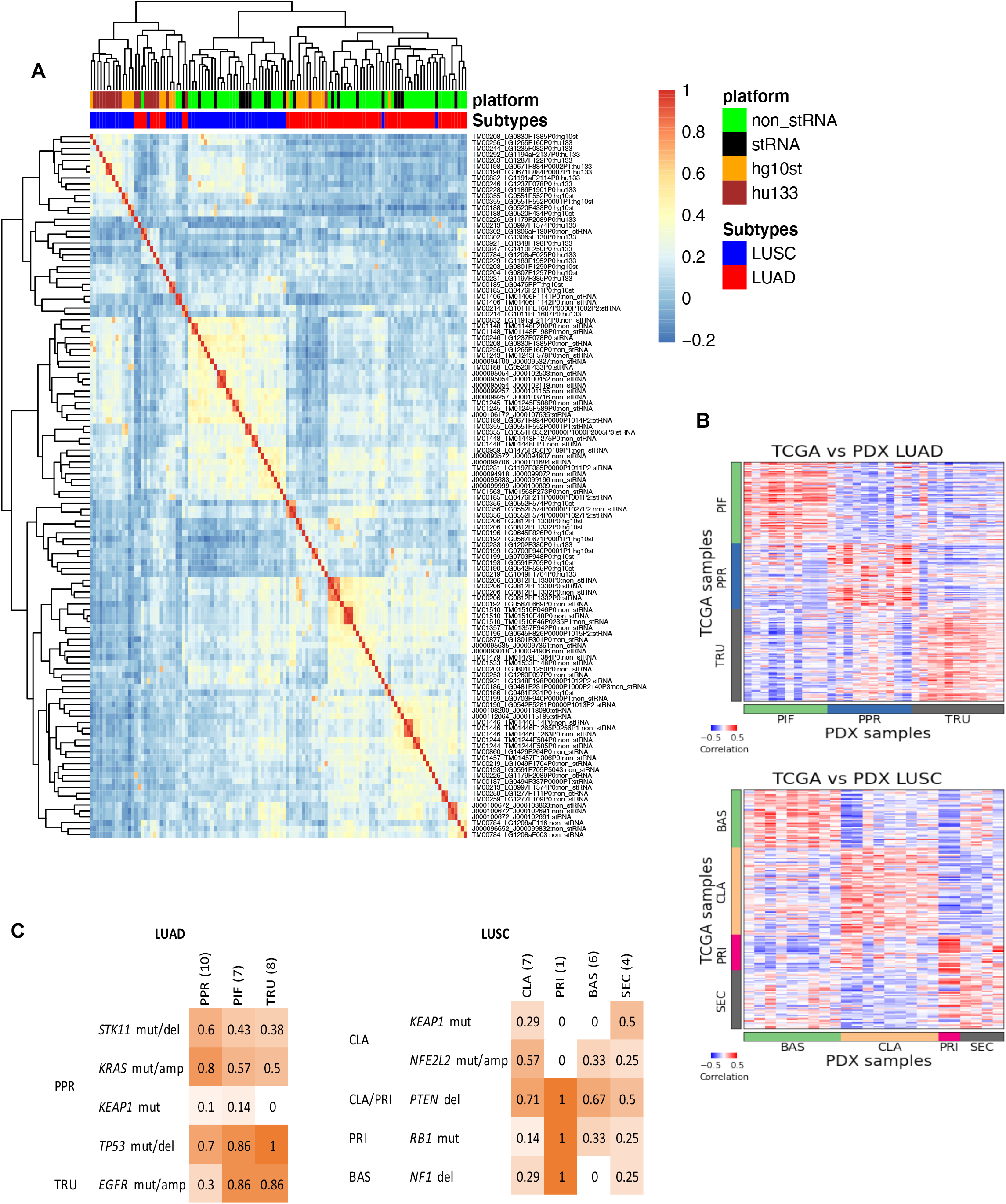
Gene expression in lung cancer PDX models. (**A)** Hierarchical clustering of gene expression percentile rank z-score for all lung cancer PDX samples and platforms. The heatmap is based on correlation values of expression percentile rank z-score expression for LUSC and LUAD PDX samples from sequencing and array platforms. The top horizontal color bars indicate the library preparation methods and platforms, and subtype designation. Sample labels are indicated by model ID, sample ID and library preparation/platform. (**B)** Expression (quantile-normalized raw RSEM counts) correlation of nearest template prediction genes between TCGA (LUAD: n=230, LUSC: n=178) and PDX (LUAD: n=36, LUSC: n=24) samples for LUAD and LUSC. The color bars indicate the subtype labels for TCGA and subtype predictions for PDX. (**C)** Proportion of PDX models with mutations reported to be enriched in LUAD and LUSC subtypes as indicated on the left. LUAD subtypes: proximal-inflammatory (PIF), proximal-proliferative (PPR), terminal respiratory unit (TRU). LUSC subtypes: basal (BAS), classical (CLA), primitive (PRI) and secretory (SEC).

It is well known that copy numbers can regulate the expression of cancer driver genes (50). We observed a positive correlation between the copy number of amplified and deleted genes and gene expression level across 63 lung PDX samples assayed for both copy number and expression profiles (Pearson correlation coefficient = 0.54, *p* < 10^−15^) for a subset of frequently amplified and deleted genes in both LUAD and LUSC PDX models (Supplementary Fig. S6A). Supplementary Fig. S6B shows the concordance between gene expression and the copy number for these genes individually. Based on these examples, it is evident that the amplification status elevates the gene expression levels of these frequently amplified genes, which are also well-known oncogenes. Similarly, the deletion status decreases the gene expression levels of these frequently deleted genes, which are also well-known TSGs.

The patient tumor associated with model TM01244 was noted as having elevated *MET* expression, but this property was only recapitulated as low-level over-expression (percentile rank z-score = 0.44, 0.51) in the engrafted tumors. Given that the *EGFR* T790M mutation present in the patient tumor was only detected by ddPCR in the PDX tumor, it is likely that the engrafted tumor is a subclone of the patient tumor where these markers are absent.

### Transcriptional Subtyping

We adapted a PDX molecular subtyping tool developed for colorectal cancer PDX models, CMScaller (32), to classify the expression subtypes for the LUAD and LUSC models (4,5,51). The previously reported gene expression subtypes for LUAD include terminal respiratory unit (TRU), proximal inflammatory (PIF) and proximal proliferative (PPR). For LUSC, the subtypes include classical (CLA), primitive (PRI), basal (BAS) and secretory (SEC) transcriptional subtypes.

For the training set, RNA-Seq data and subtype labels for LUAD and LUSC from TCGA were used (7,33). This analysis yielded 793 and 1224 template genes to classify LUAD and LUSC subtypes respectively (Supplementary Table S7 and S8), resulting in high accuracies of 93% and 92% for the TCGA validation set (Supplementary Table S4). For the JAX PDX models, 31 out of 36 LUAD samples and 24 out of 24 LUSC samples were classified in expression subtype categories with high confidence (FDR-adjusted P-values < 0.05) (Supplementary Tables S5 and S6). Among the LUAD samples, 32% were classified as PIF, 32% as PPR and 35% as TRU. Among the LUSC samples, 38% were classified as BAS, 38% were classified as CLA, 8% were classified as PRI, and 17% were classified as SEC. For models with multiple samples, all were predicted as the same subtype within each model, except for LUSC model TM01448 in which PT and P0 were classified as BAS and SEC, respectively. Spatial tumor heterogeneity in the tumor sample used to establish the PDX model is a plausible explanation for these classification differences (52). Indeed, the patient and P0 tumor samples share the same clinically relevant mutations except for a PTEN nonsense mutation that was detected only in the PT sample.

To further confirm the reliability of the classifications, we compared the expression of the template genes between TCGA samples with known subtype labels and the predicted subtypes of the samples of the PDX models. We observed high correlation within the template genes of each respective LUAD or LUSC subtype (Fig. 4B). This confirms that the expression level of the template genes is replicated in the lung cancer PT/PDX samples. The subtypes were also enriched in other genome alteration profiles (mutations and copy number aberrations) (5,7,33,51). Despite the limited number of samples, we observed higher proportion in some of the reported subtype-enriched alterations within the respective predicted PDX subtypes (Fig. 4C). In particular, the LUAD PPR subtype was reported to be enriched in *STK11* and *KRAS* alterations in other PT datasets, and the PDX models classified as PPR subtype showed higher frequencies of these alterations compared other subtypes. The same observation can also be made for the *NFE2L2* alteration enriched in LUSC CLA subtype. As such, the PDX models displayed subtype-specific expression and/or alteration profiles similar to those reported in patient tumor subtyping studies.

### Treatment responses in PDX models

The lung cancer PDX models in the JAX repository originated from both treatment-naïve and previously treated patient tumors. Many of the tumors submitted for PDX generation were selected based on the presence of clinically relevant mutations per National Comprehensive Cancer Network (NCCN) guidelines (Supplementary Table S1) (10,25,53). Eighteen of the models have tumors that harbor activating mutations in the KRAS gene, of increased clinical significance due to the recent development of small molecule inhibitors to *KRAS* G12C-mutated cancers. Cohorts of tumor-bearing mice of a subset of the lung cancer PDX models were enrolled in dosing studies to evaluate responses to drug treatment.

#### Targeted treatment of EGFR mutant PDXs

Nine PDX models in the JAX collection harbor activating mutations in *EGFR* (L858R, exon 19 deletion, exon 20 insertion) and were tested for response to tyrosine kinase inhibition. Six of the models (TM00199, TM00204, TM00219, TM00253, and TM00784) were derived from patients at the time of progression on either single-agent or combinations of erlotinib; two of the models (J000100672 and TM00193) were derived from treatment-naïve patients. Both TM00204 and TM00219 harbor the *EGFR* T790M mutation, and TM00784 harbors *MET* amplification. These markers are associated with acquired resistance to treatment with tyrosine kinase inhibitors (10,53). J000100672 harbors the exon 20 insertion associated with *de novo* resistance to TKI inhibitors (54). TM00253 harbors the mutation *EGFR* V834L, which is associated with decreased response to erlotinib (55). Cohorts of tumor-bearing mice of these models, except for TM00193, were treated with single-agent erlotinib. TM00199 displayed partial response, TM00253 displayed stable disease, and the other four models with TKI treatment resistance mutations displayed progressive disease (Fig. 5A, Supplementary Fig. S7 and S8). The overall lack of complete response recapitulated the treatment response observed in the patients and the response expected from the *EGFR* mutation status of these models.

**Figure 5.**
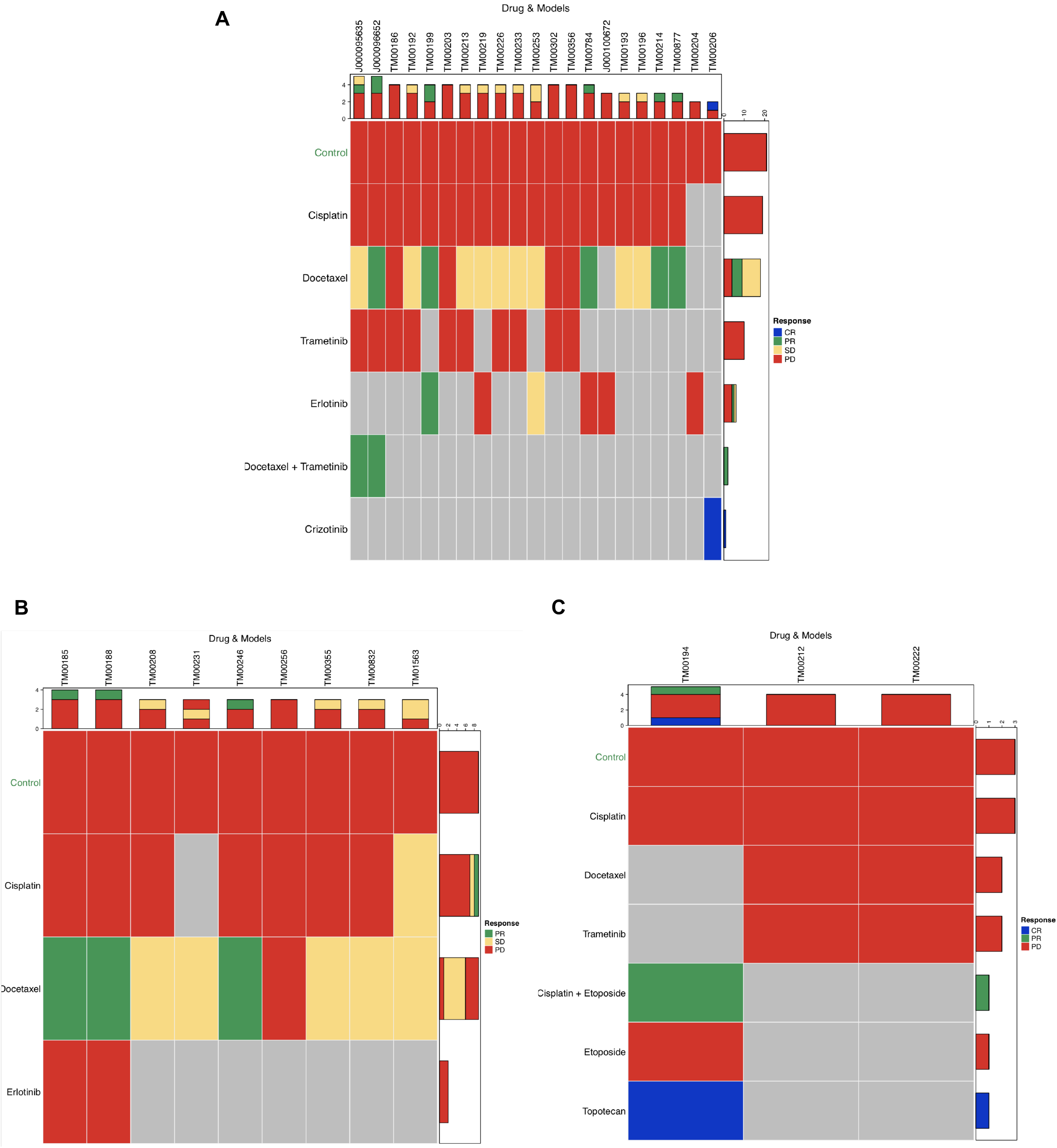
Treatment response of cancer drugs on the PDX models **A,B,C**. Modified RECIST classification summary plot for PDX models classified as **(A)** LUAD, **(B)** LUSC, and **(C)** all other lung cancer types. Colors indicate RECIST classification for a treatment cohort (CR: complete response, PR: partial response, SD: stable disease, and PD: progressive disease). Number of treatments in each RECIST category is shown at the top, and number of models in each RECIST category is shown on the right side of each plot. Plots were generated with the R package Xeva (version 1.6.0).

Two models (TM00199 and TM00219) were tested using second-generation treatment strategies to test treatment options following the development of resistance to first-generation TKIs (56). The cohort of tumor bearing animals from these models received the combination of afatinib and cetuximab along with single-agent erlotinib, afatinib, cetuximab, and vehicle controls for a 21-day study followed by an observation period of up to 90 days. As expected, TM00199 did not respond to treatment with erlotinib. Single-agent afatinib-treated animals subsequently progressed after a succession of treatment, whereas animals treated with cetuximab (with or without afatinib) exhibited complete response. The treatment effects on *EGFR* expression and phosphorylation were examined at six and 24-hour timepoints (Figure 6). After a single treatment, the *EGFR* tyrosine kinase inhibitors erlotinib and afatinib induced near-complete downregulation of *EGFR* phosphorylation within six hours, rebounding to control levels after 24 hours (Figure 6A). In contrast, cetuximab showed moderate downregulation by six hours and complete downregulation at 24 hours accompanied by diminished total protein expression. The combination of afatinib plus cetuximab resulted in ablated phosphorylation at six hours, maintained at the 24 timepoints, associated with reduced protein expression.

**Figure 6.**
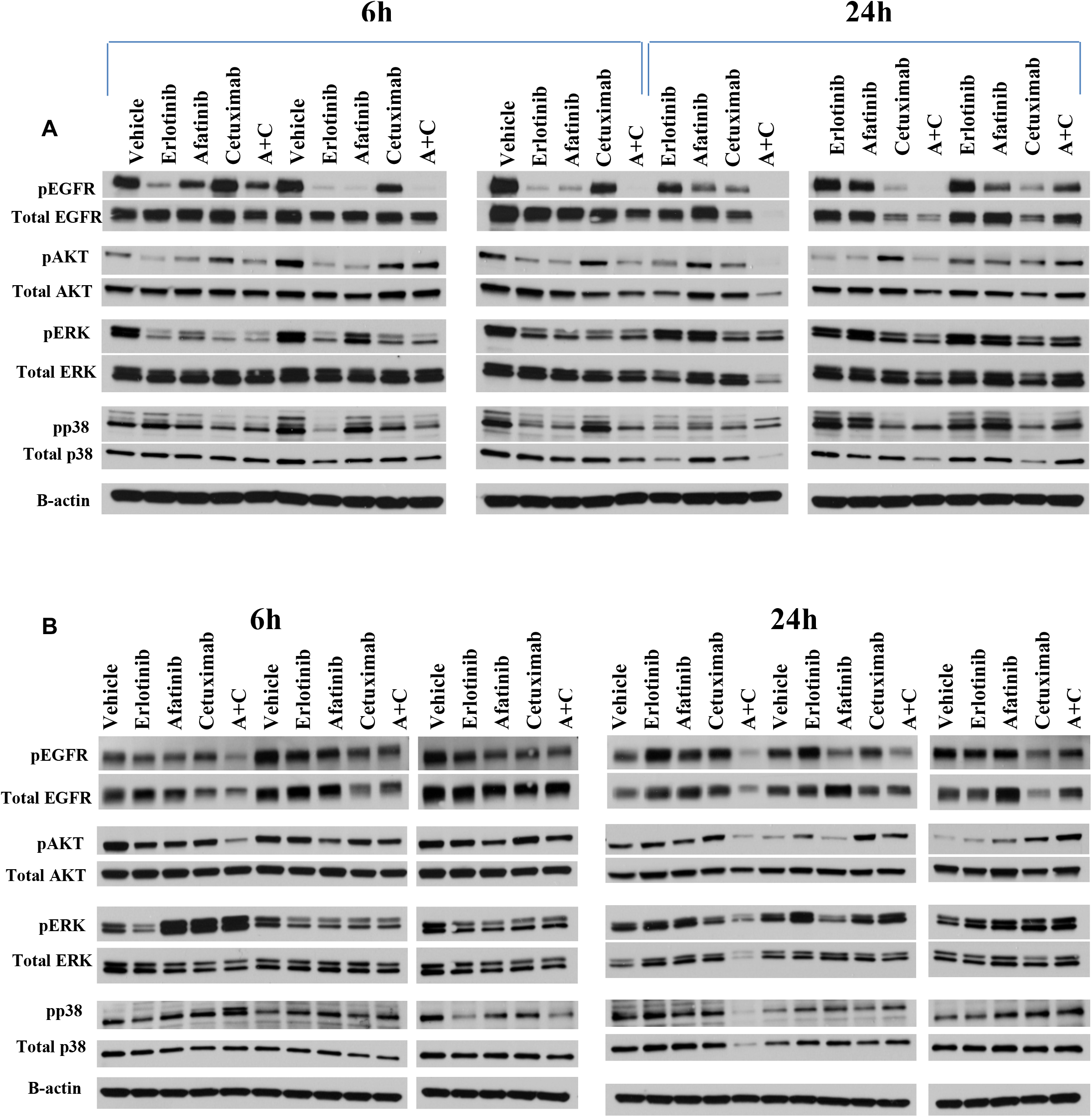
Treatment induced changes in phosphorylation of EGFR, AKT, ERK, P38 at 6 and 24 hours in lung PDX models treated with erlotinib, afatinib, cetuximab, and afatinib+cetuximab. **(A)** In model TM00199, a single treatment of erlotinib and afatinib induced near-complete downregulation of *EGFR* phosphorylation within six hours, rebounding to control levels after 24 hours. Treatment with cetuximab demonstrated moderate downregulation by six hours and complete downregulation at 24 hours accompanied by diminished total protein expression. The combination of afatinib plus cetuximab resulted in ablated phosphorylation at six hours, maintained at the 24 timepoints, associated with reduced protein expression. (**B**) In model TM00219 none of the EGFR-targeted agents could entirely suppress *EGFR* phosphorylation at 6 or 24 hours.

The TM00219 model was derived from a patient at the time of erlotinib progression, associated with the emergence of the T790M *EGFR* resistance mutation which was observed in both the patient post-erlotinib treatment biopsy and the engrafted tumor. This model showed no benefit from erlotinib or cetuximab, with only marginal albeit statistically significant activity from afatinib. In this unresponsive model, none of the EGFR-targeted agents could entirely suppress *EGFR* phosphorylation at six or 24 hours (Figure 6B).

#### Targeted treatment of an EML4-ALK fusion PDX model

LUAD model TM00206 harbors the *EML4-ALK* fusion and, as expected, had a robust response to treatment with the ALK tyrosine kinase inhibitor, crizotinib (Fig. 5A, Supplementary Fig. S7 and S9) (57). The response at the cohort level was categorized as a complete response (CR). However, 2 of the 9 mice in the treatment cohort were classified as partial response (PR). Acquired crizotinib resistance has been reported in *ALK*-rearranged NSCLCs (58) and the clinical records for the patient reveal that the individual’s cancer progressed while on treatment. The variability in the response in the corresponding PDX models may be due to the presence of resistant subclones. Although the treated PDXs were not tested for known resistance variants, the genomic data from two early passage (P0) tumors for this model revealed the presence of reported resistance mutations at a subclonal level. The *ALK* L1196M mutation was detected at an allele frequency of 22% and a low-level *KIT* amplification (log(CN/ploidy) = 0.43) was detected in one of the P0 tumors (LG0812PE1330P0), while low-level amplification in *ALK* and *EML4*, possibly the fusion, was detected in another P0 sample (LG0812PE1332P0).

#### Treatment of KRAS-mutant PDXs

Twelve models harboring various gain-of-function *KRAS* mutations (Supplementary Table 1) were treated with a *MEK1/2* inhibitor (trametinib), which acts through the inhibition of the MAPK pathway downstream of *KRAS*. The treatment responses were classified as progressive disease for all models which is consistent with the results in clinical trials where single-agent trametinib showed no improved efficacy compared to docetaxel in patients with advanced *KRAS*-mutant NSCLC (Fig. 5, Supplementary Fig. S7) (59). Partial response was observed for the combination of docetaxel and trametinib for *KRAS*-mutant (G12D) model J000095635 compared to single-agents (stable and progressive disease, respectively). For model J000096652 (*KRAS* G12C), the combination docetaxel and trametinib showed no additional benefit over single-agent docetaxel. Both treatment arms were classified as PR (Fig. 5A, Supplementary Fig. S7 and S10).

## Discussion

The treatment of NSCLC has rapidly evolved, with chemotherapy being replaced by either targeted therapies or immunotherapy in many clinical scenarios. In order for continued advances to be made, preclinical models which accurately reflect the complexity and heterogeneity of human cancers, as well as being predictive of drug sensitivity and resistance patterns observed in patients, are mandatory. Given the complexities of drug-tumor interactions together with inter- and intra-patient tumor heterogeneity, PDX models stand apart in the preclinical arena as experimentally tractable and reproducible models that recapitulate the clinically relevant genomic properties and treatment responses of the patient tumors from which they are derived.

The repository of lung cancer PDX models maintained at The Jackson Laboratory was generated, characterized, and annotated in collaboration with clinical investigators. For models with corresponding patient tumor genomic data, the implanted tumors in the lung PDXs maintained the histological characteristics and genomic properties of patient tumors from which they were derived. Treatment responses for targeted agents in the models were consistent with expectations based on the presence of specific molecular targets and also recapitulated the clinical outcomes of patients who subsequently received the same therapies. The PDX models available from the JAX repository are a validated resource for preclinical investigations into the efficacy of new cancer treatments and for basic research into the mechanisms of acquired resistance to target-directed therapies and for developing strategies to overcome treatment resistance.

## Acknowledgements

The authors express their deep appreciation to the patients who consented to have their tumor tissues used to generate the patient-derived cancer models described in this manuscript to make these models broadly available for basic and pre-clinical cancer research. The generation and characterization of models in the JAX lung cancer PDX repository was funded by The Jackson Laboratory Director’s Innovation Fund, the Maine Cancer Foundation, the Kleeberg Foundation, the Hope Foundation, and the Integrated Translational Science Center (ITSC) (5U01 CA180944). Genome data analysis was supported, in part, by the JAX Cancer Center Computational Sciences Shared Service (P30 CA034196). The PDX Web portal was developed with support from R01 CA089713. Additional support provided by Susan and Jerry Knapp and the Addario Foundation and Stand Up to Cancer (201502309). The UCD Pathology Biorepository assisted with collection and histological interpretation of specimens. The content is solely the responsibility of the authors and does not necessarily represent the official views of the NIH.

