## Supplemental Figure 1 for "A Genomically and Clinically Annotated Patient Derived Xenograft (PDX) Resource for Preclinical Research in Non-Small Cell Lung Cancer"

**Supplementary Figure S1.** Percentage of human and mouse reads identified by Xenome for CTP and RNA-Seq data of all lung cancer PDX samples.

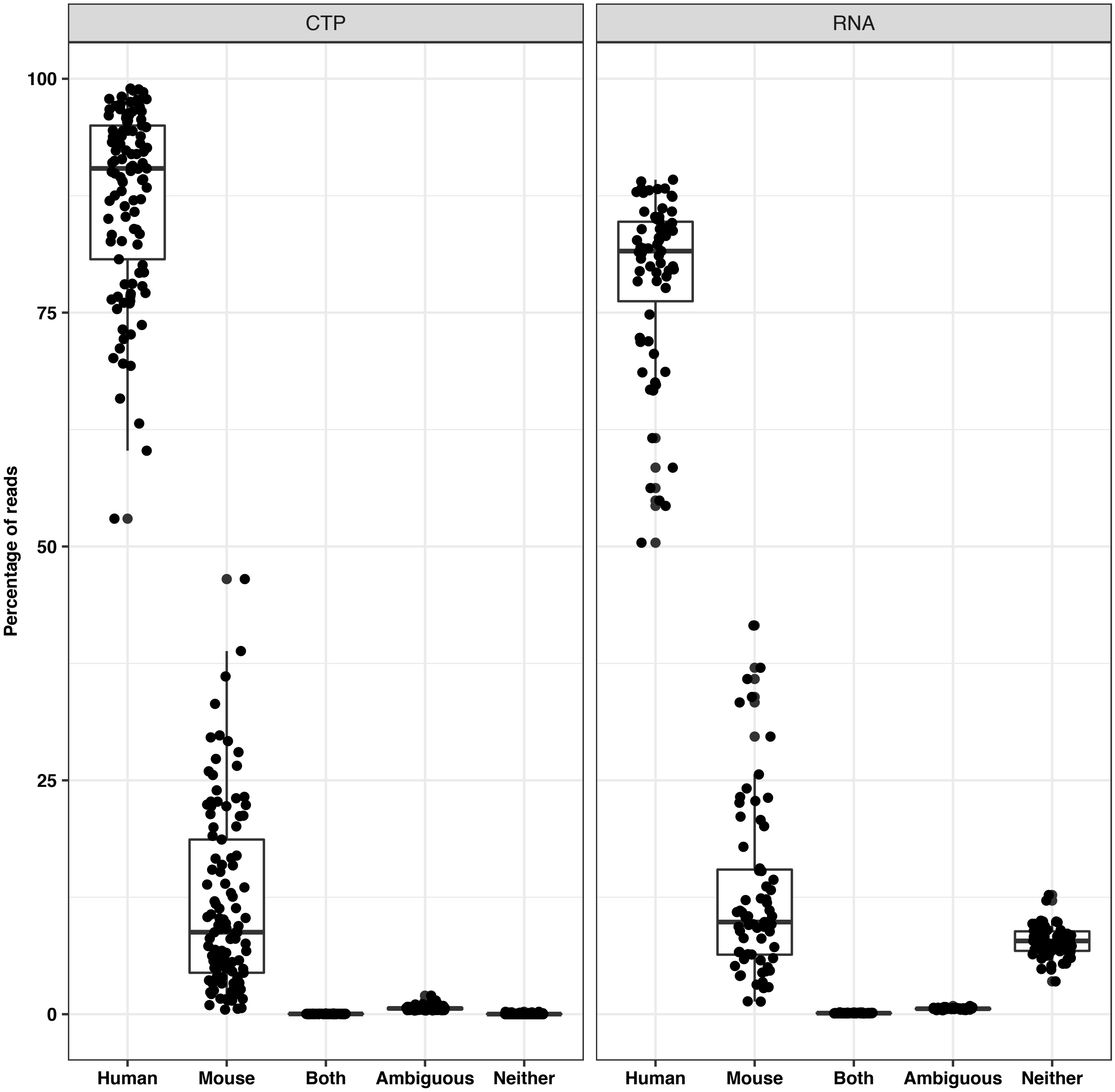
