## Supplemental Figure 2 for "A Genomically and Clinically Annotated Patient Derived Xenograft (PDX) Resource for Preclinical Research in Non-Small Cell Lung Cancer"

**Supplementary Figure S2.** Percentage of tumor stages in the JAX PDX and TCGA PanCancer Atlas LUAD and LUSC datasets.

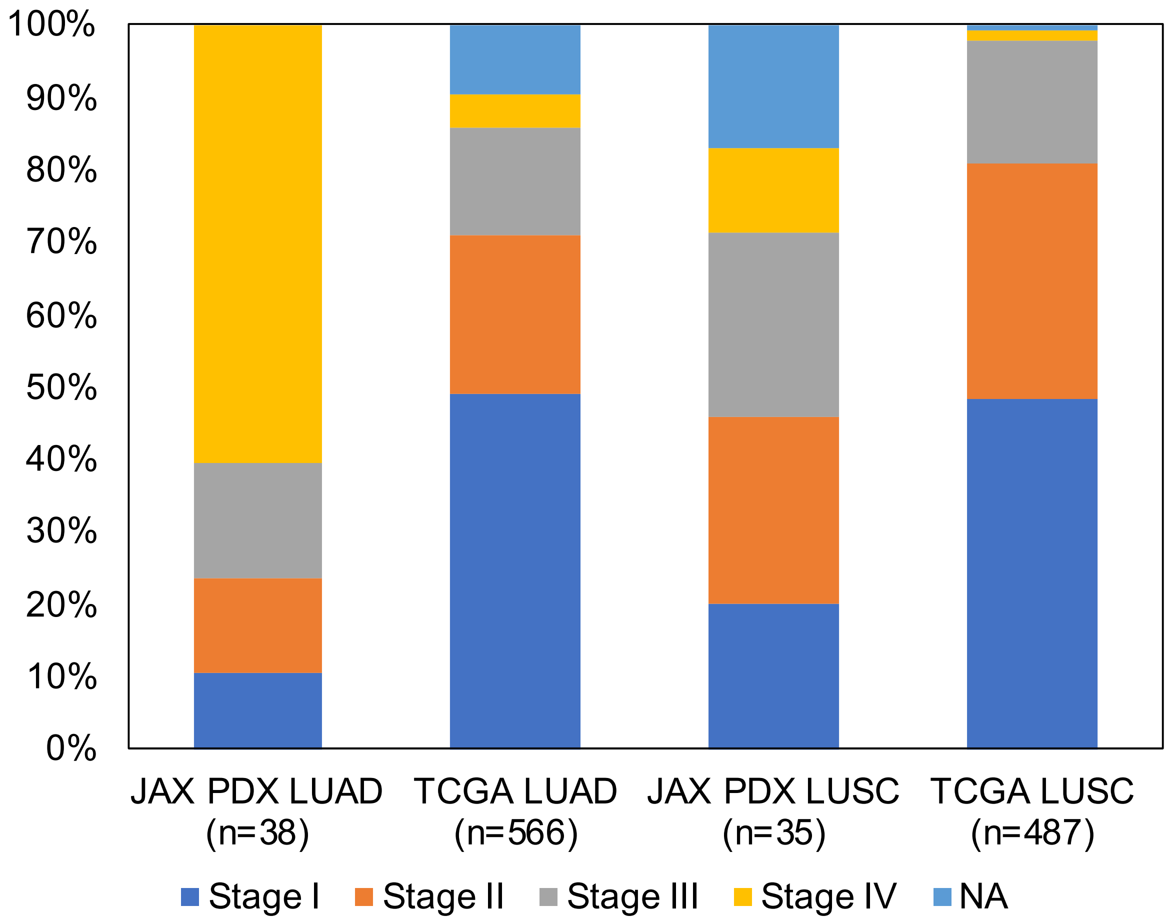
