## Supplemental Figure 3 for "A Genomically and Clinically Annotated Patient Derived Xenograft (PDX) Resource for Preclinical Research in Non-Small Cell Lung Cancer"

**Supplementary Figure S3.** Distribution of TMB and MSI scores in LUAD and LUSC PDX samples based on CTP sequencing data.

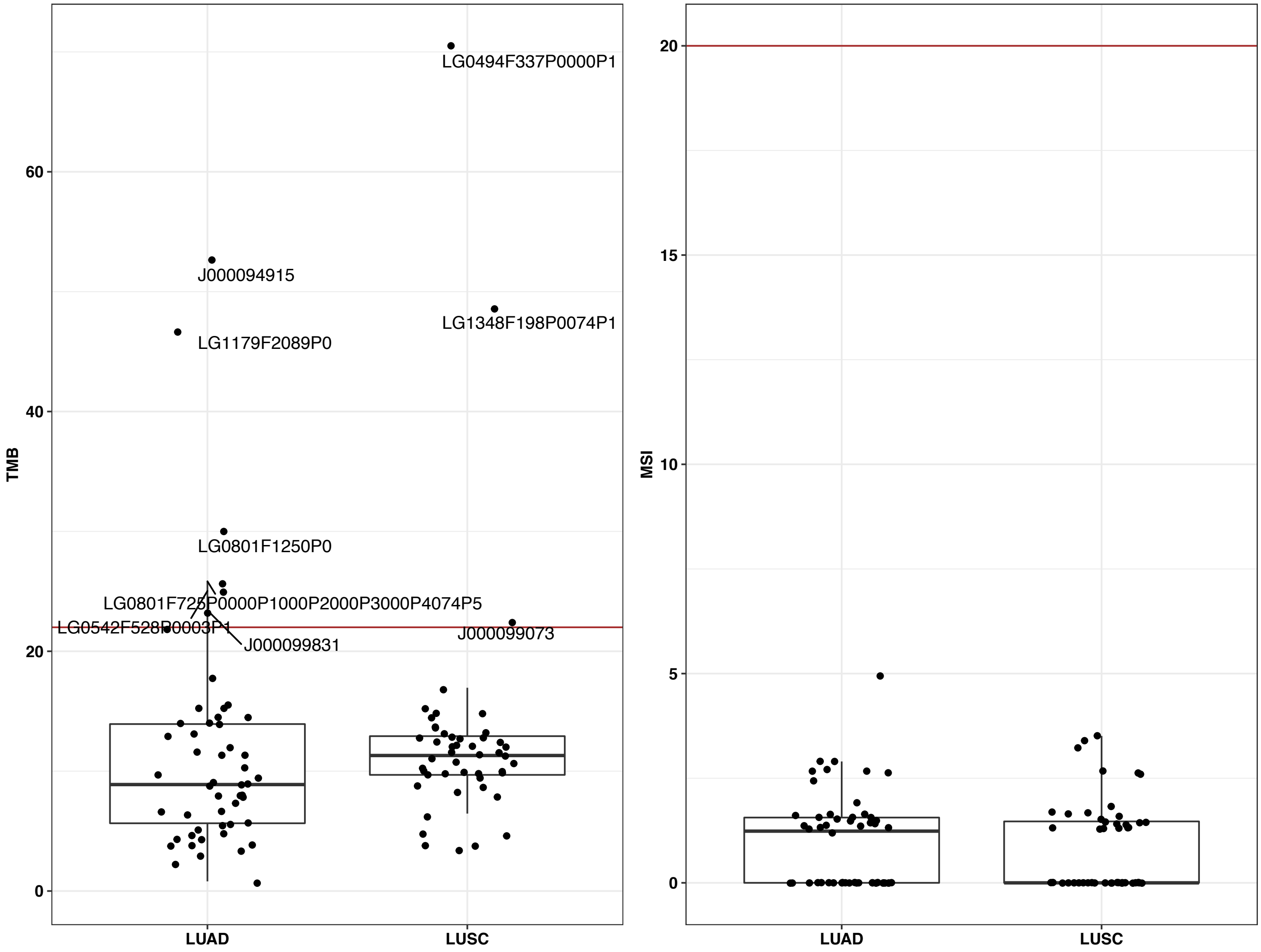
