## Supplemental Figure 4 for "A Genomically and Clinically Annotated Patient Derived Xenograft (PDX) Resource for Preclinical Research in Non-Small Cell Lung Cancer"

**Supplementary Figure S4.** Copy number profile profiles of PDX samples classified as **(A)** LUAD, **(B)** LUSC, and **(C)** other subtypes (adenosquamous cell carcinoma (TM01031), large cell neuroendocrine carcinoma (TM00212, TM00694), and small cell carcinoma (J00093933, J000112882, TM00194, TM01284)).

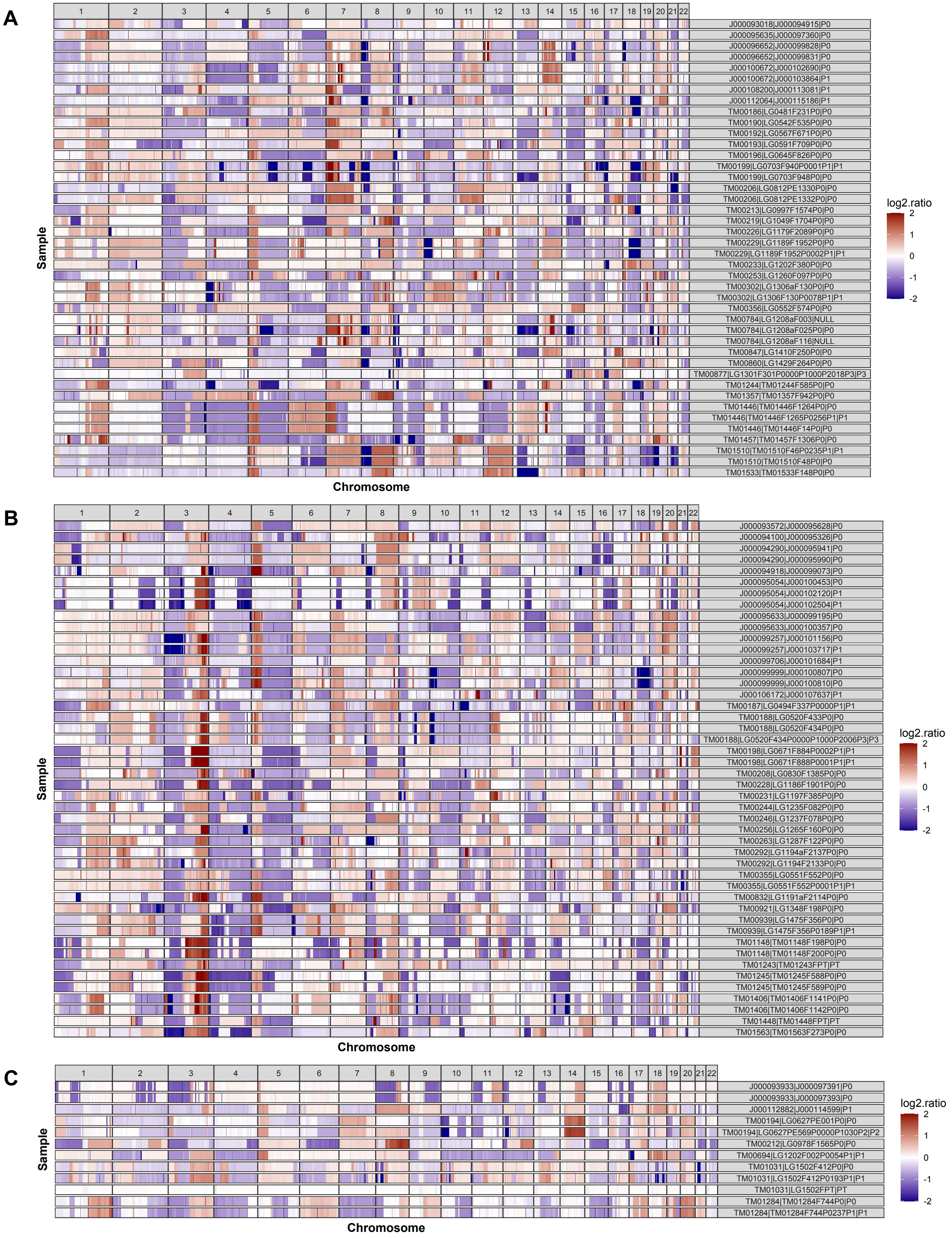
