## Supplemental Figure 5 for "A Genomically and Clinically Annotated Patient Derived Xenograft (PDX) Resource for Preclinical Research in Non-Small Cell Lung Cancer"

**Supplementary Figure S5.** Distribution of expression correlation values in Fig. 4A, between different PDX models, same models assayed on the same platform and library preparation method, and same models assayed on different platforms and library preparation methods.

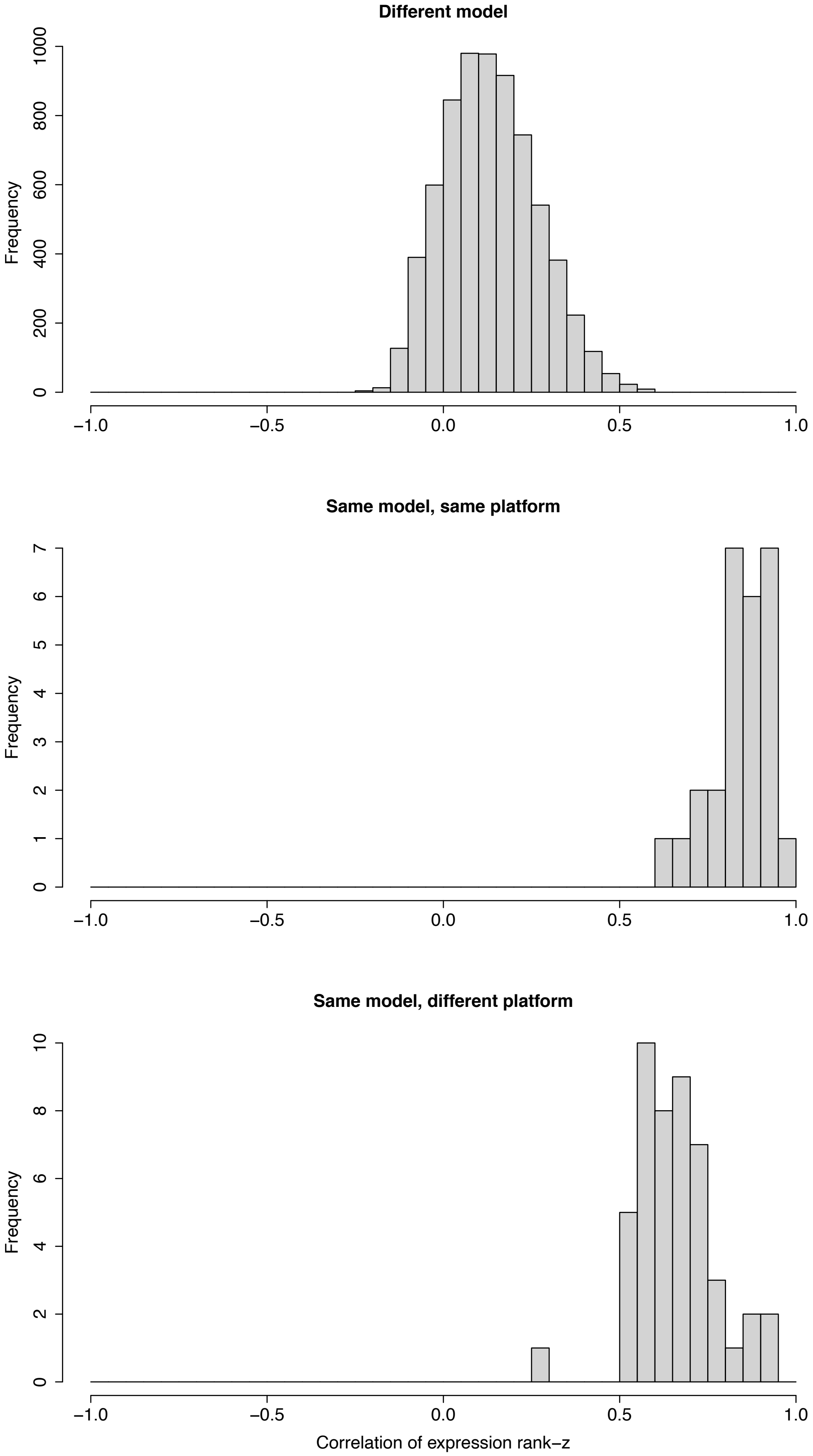
