## Supplemental Figure 6 for "A Genomically and Clinically Annotated Patient Derived Xenograft (PDX) Resource for Preclinical Research in Non-Small Cell Lung Cancer"

**Supplementary Figure S6.** A. Correlation of copy number and gene expression for amplified ($log_2$(CN ratio) > 1) and deleted ($log_2$(CN ratio) < -1) selected genes in PDX samples assayed for both copy number and expression profiles. B. Effect of copy number on gene expression percentile rank z-score of selected genes in PDX samples. EGFR, MYC and KRAS are frequently amplified genes in LUAD and LUSC. APC, CDKN2A and TP53 are frequently deleted genes in LUAD and LUSC.

A

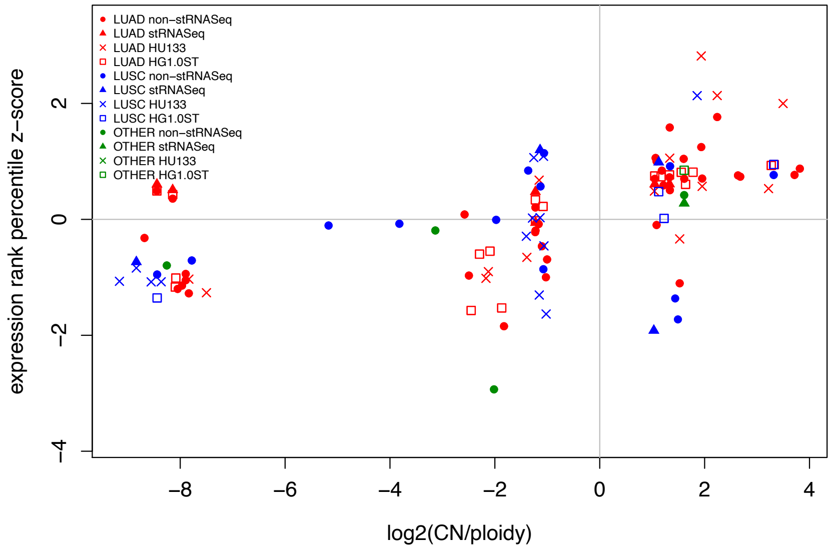


B

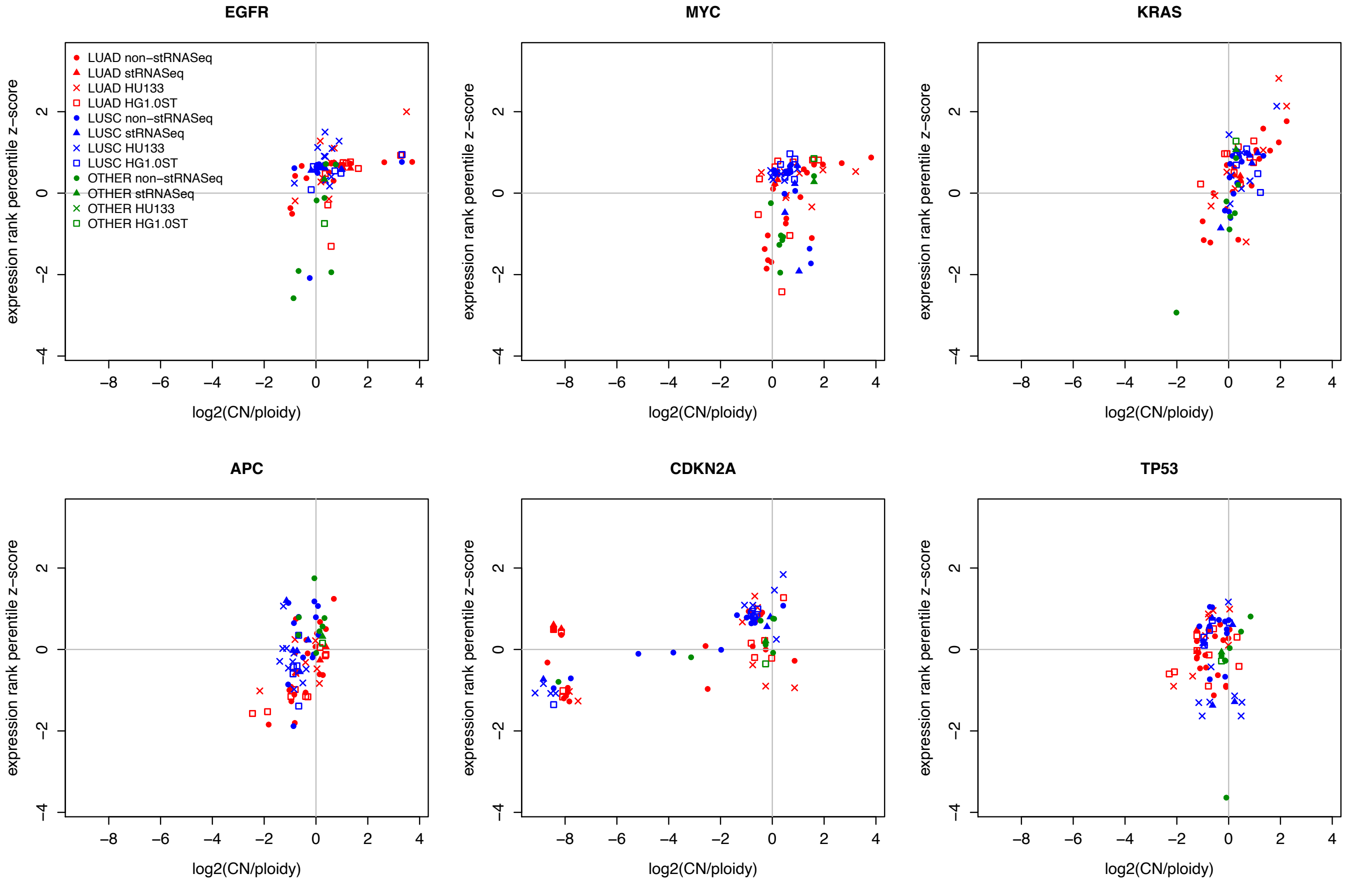
