## Supplemental Figure 7 for "A Genomically and Clinically Annotated Patient Derived Xenograft (PDX) Resource for Preclinical Research in Non-Small Cell Lung Cancer"

**Supplementary Figure S7.** Treatment response of cancer drugs on the PDX models per mouse corresponding to Fig 5. Modified RECIST classification summary plot for PDX models classified as **(A)** LUAD, **(B)** LUSC, and **(C)** all others. Colors indicate RECIST classification for a treatment cohort (CR: complete response, PR: partial response, SD: stable disease, and PD: progressive disease). Number of treatments in each RECIST category is shown at the top, and number of mouse in each RECIST category is shown on the right side of each plot. These plots were generated with the R package Xeva (version 1.6.0).

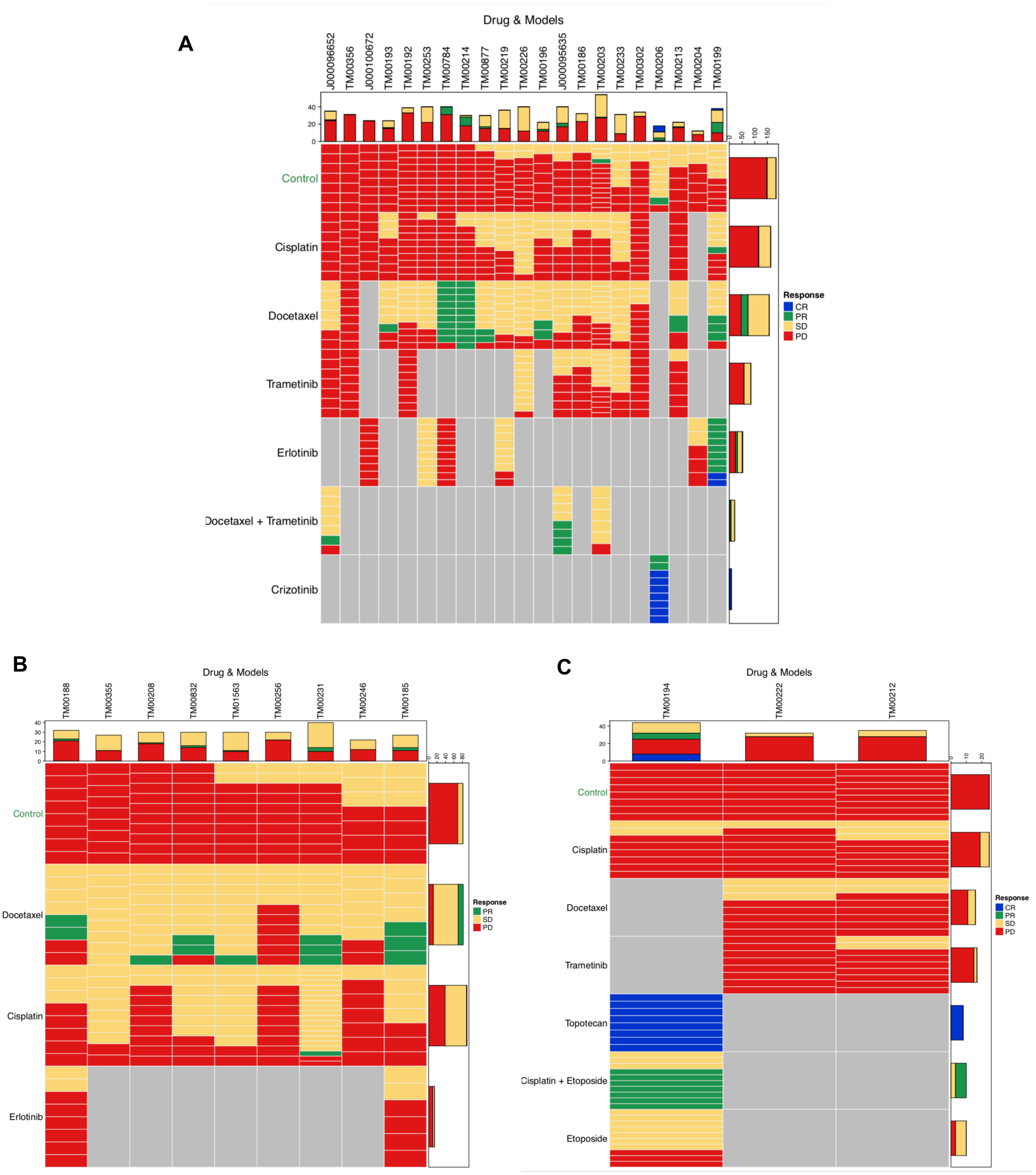
