## Supplemental Figure 8 for "A Genomically and Clinically Annotated Patient Derived Xenograft (PDX) Resource for Preclinical Research in Non-Small Cell Lung Cancer"

**Supplementary Figure S8.** Treatment response results for LUAD PDX models TM00199, TM00204, TM00219, TM00253, TM00784 and J000100672 illustrated in absolute tumor volume, change in tumor volume and tumor growth inhibition. None of these models display complete response to erlotinib treatment.

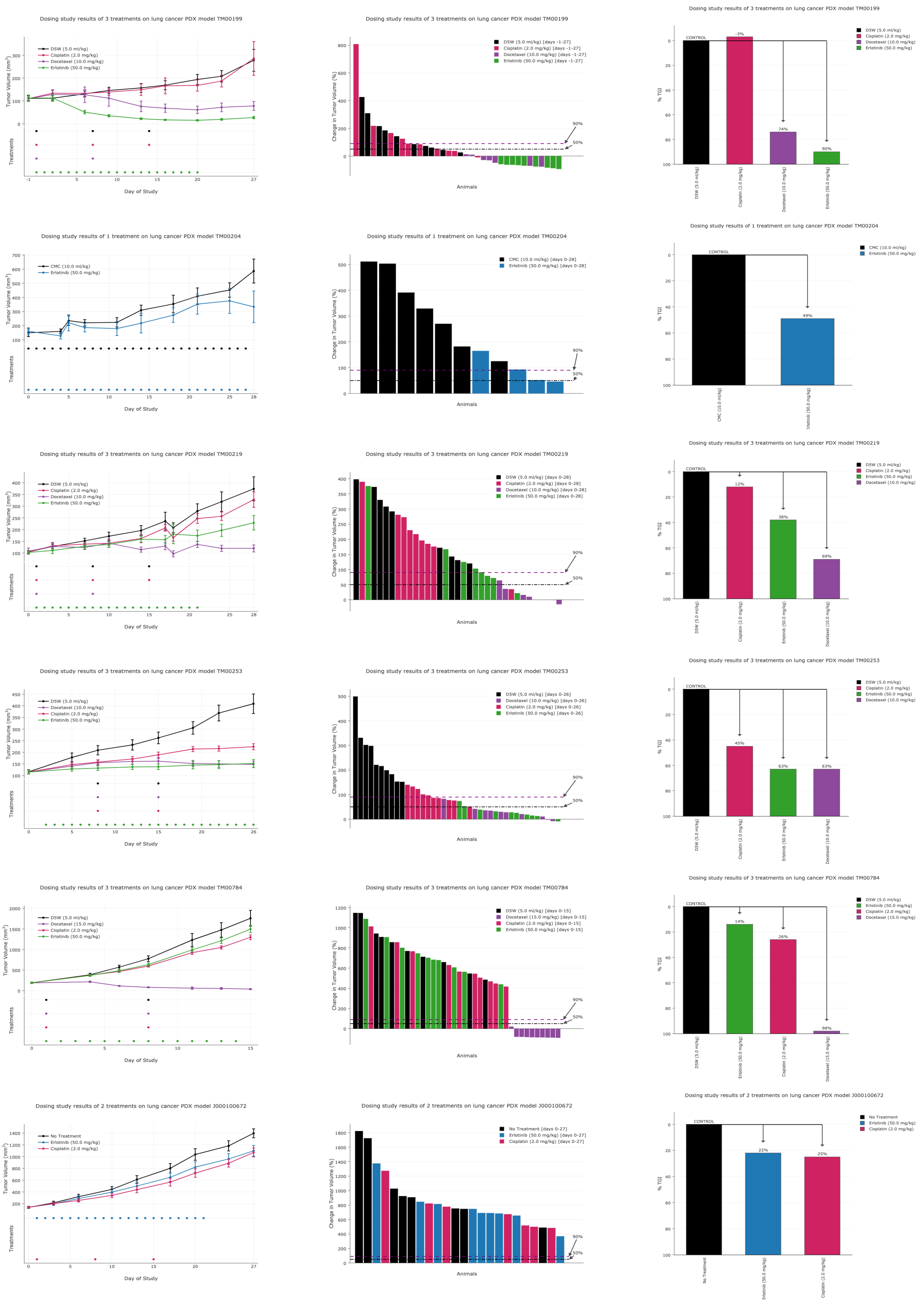
