## Supplemental Figure 9 for "A Genomically and Clinically Annotated Patient Derived Xenograft (PDX) Resource for Preclinical Research in Non-Small Cell Lung Cancer"

**Supplementary Figure S9.** Treatment response results for LUAD PDX model T00206 illustrated in absolute tumor volume, change in tumor volume and tumor growth inhibition. The model shows complete response to crizotonib treatment but two of the animals show less than 50% reduction in tumor volume.

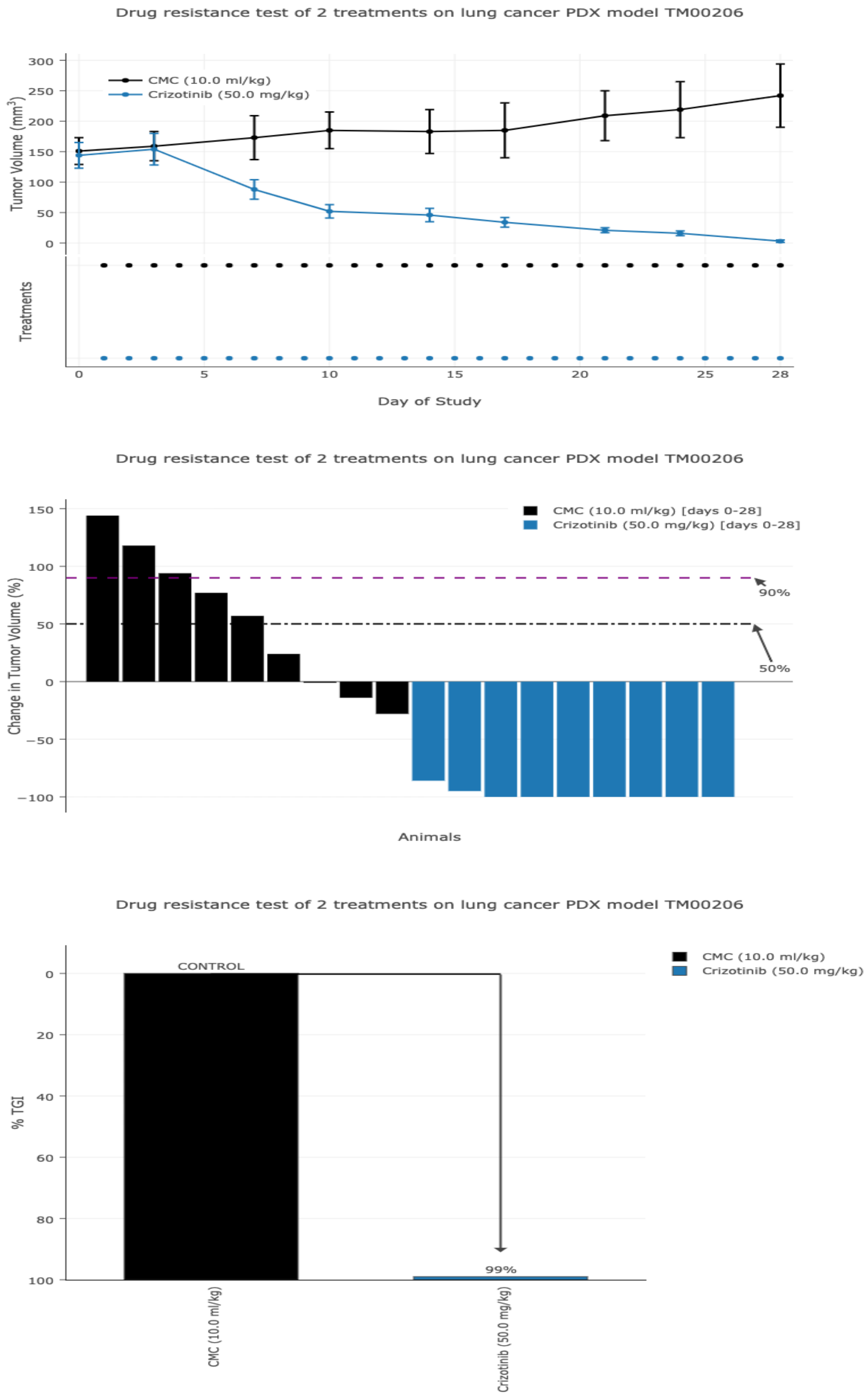
