## Supplemental Figure 10 for "A Genomically and Clinically Annotated Patient Derived Xenograft (PDX) Resource for Preclinical Research in Non-Small Cell Lung Cancer"

**Supplementary Figure S10.** Treatment response results for LUAD models J000095635 and J000096652 illustrated in absolute tumor volume, change in tumor volume and tumor growth inhibition. Both models display partial response to combination treatment of docetaxel and trametinib.

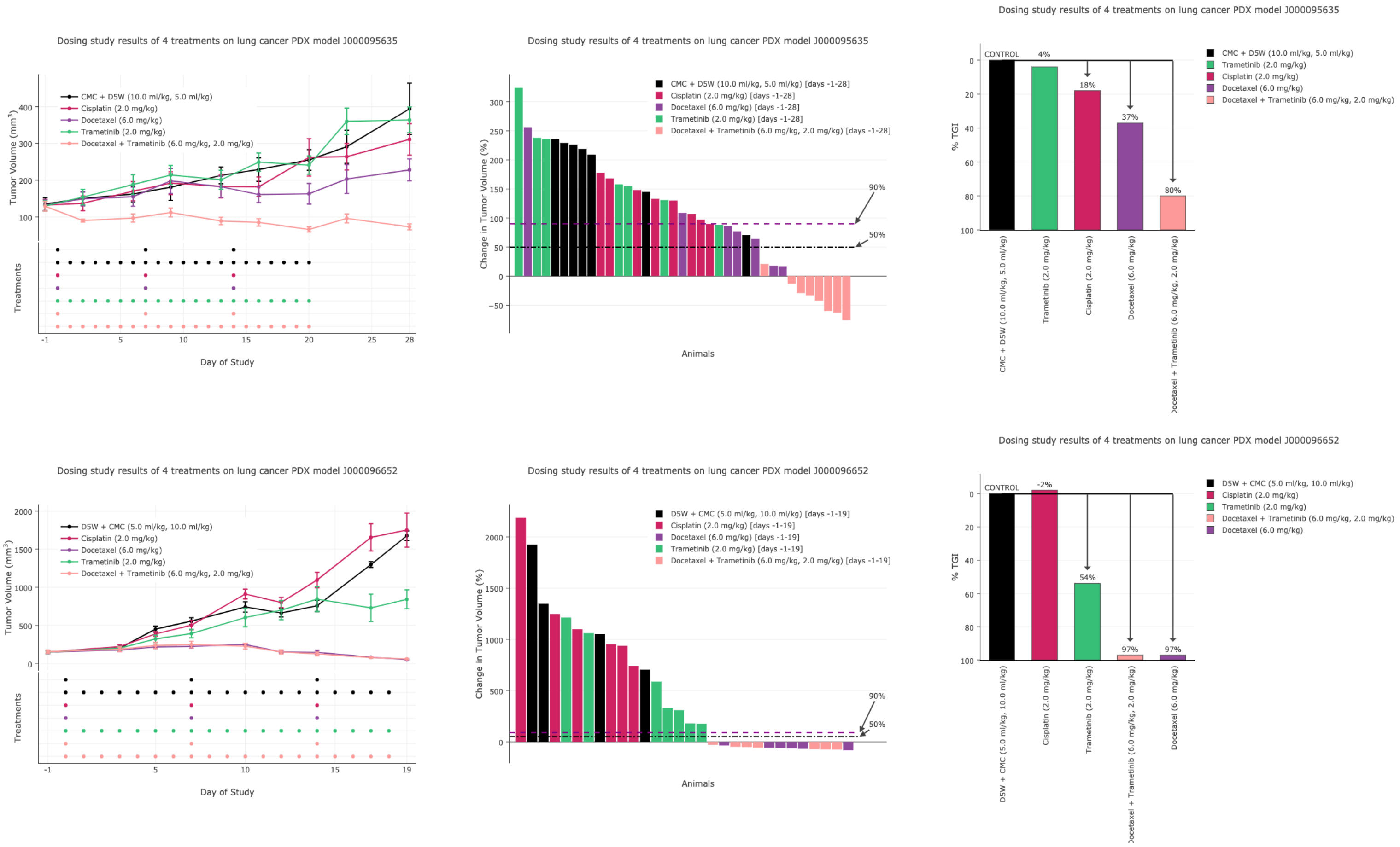
