## Supplemental Table 4 for "A Genomically and Clinically Annotated Patient Derived Xenograft (PDX) Resource for Preclinical Research in Non-Small Cell Lung Cancer"

**Supplementary Table S4.** Accuracies of expression subtyping using nearest template prediction of TCGA LUAD and LUSC RNA-Seq data.

| LUAD: 3,547 filtered genes | | | | |
|---|---|---|---|---|
| **Subtypes (Template genes)** | **Precision** | **Recall** | **F1-score** | **Support** |
| PIF (232) | 0.86 | 0.92 | 0.89 | 13 |
| PPR (264) | 1.00 | 0.83 | 0.91 | 12 |
| TRU (297) | 0.94 | 1.00 | 0.97 | 17 |
| **accuracy** | | | 0.93 | 42 |
| **macro avg** | 0.93 | 0.92 | 0.92 | 42 |
| **weighted avg** | 0.93 | 0.93 | 0.93 | 42 |

| LUSC: 3,564 filtered genes | | | | |
|---|---|---|---|---|
| **Subtypes (Template genes)** | **Precision** | **Recall** | **F1-score** | **Support** |
| BAS (294) | 1.00 | 0.89 | 0.94 | 9 |
| CLA (361) | 0.87 | 1.00 | 0.93 | 13 |
| PRI (281) | 1.00 | 0.80 | 0.89 | 5 |
| SEC (288) | 0.89 | 0.89 | 0.89 | 9 |
| **accuracy** | | | 0.92 | 36 |
| **macro avg** | 0.94 | 0.89 | 0.91 | 36 |
| **weighted avg** | 0.92 | 0.92 | 0.92 | 36 |
