## Supplemental Table 5 for "A Genomically and Clinically Annotated Patient Derived Xenograft (PDX) Resource for Preclinical Research in Non-Small Cell Lung Cancer"

**Supplementary Table S5.** Predicted LUAD subtypes of JAX PDX models.

| Sample name | Model name | Subtype | FDR |
|---|---|---|---|
| J000094906 | J000093018 | PPR | 0.016 |
| J000097361 | J000095635 | PPR | 0.001 |
| J000099832 | J000096652 | PIF | 0.001 |
| J000102691 | J000100672 | PIF | 0.756 |
| J000103863 | J000100672 | PIF | 0.022 |
| LG0481F231P0000P1000P2140P3 | TM00186 | TRU | 0.001 |
| LG0567F669P0 | TM00192 | TRU | 0.003 |
| LG0591F705P5043 | TM00193 | TRU | 0.001 |
| LG0703F940P0000P1 | TM00199 | TRU | 0.001 |
| LG0801F1250P0 | TM00203 | PPR | 0.001 |
| LG0812PE1330P0 | TM00206 | PIF | 0.001 |
| LG0812PE1332P0 | TM00206 | PIF | 0.069 |
| LG0997F1574P0 | TM00213 | PPR | 0.001 |
| LG1049F1704P0 | TM00219 | TRU | 0.001 |
| LG1179F2089P0 | TM00226 | TRU | 0.274 |
| LG1260F097P0 | TM00253 | PIF | 0.003 |
| LG1277F109P0 | TM00259 | TRU | 0.001 |
| LG1277F111P0 | TM00259 | TRU | 0.001 |
| LG1306aF130P0 | TM00302 | PIF | 0.001 |
| LG0552F574P0000P1027P2 | TM00356 | PPR | 0.001 |
| LG1208aF003 | TM00784 | PIF | 0.001 |
| LG1208aF116 | TM00784 | PIF | 0.001 |
| LG1429F264P0 | TM00860 | PPR | 0.001 |
| LG1301F301P0 | TM00877 | PIF | 0.076 |
| TM01244F584P0 | TM01244 | TRU | 0.001 |
| TM01244F585P0 | TM01244 | TRU | 0.001 |
| TM01357F942P0 | TM01357 | PPR | 0.001 |
| TM01446F1263P0 | TM01446 | TRU | 0.368 |
| TM01446F1265P0256P1 | TM01446 | TRU | 0.026 |
| TM01446F14P0 | TM01446 | TRU | 0.001 |
| TM01457F1306P0 | TM01457 | PPR | 0.001 |
| TM01479F1384P0 | TM01479 | PPR | 0.001 |
| TM01510F046P0 | TM01510 | PIF | 0.001 |
| TM01510F46P0235P1 | TM01510 | PIF | 0.001 |
| TM01510F48P0 | TM01510 | PIF | 0.001 |
| TM01533F148P0 | TM01533 | PPR | 0.001 |
