## Supplemental Table 6 for "A Genomically and Clinically Annotated Patient Derived Xenograft (PDX) Resource for Preclinical Research in Non-Small Cell Lung Cancer"

**Supplementary Table S6. Predicted LUSC subtypes of JAX PDX models.**

| Sample name | Model name | Subtype | FDR |
|---|---|---|---|
| J000094937 | J000093572 | SEC | 0.001 |
| J000095327 | J000094100 | CLA | 0.001 |
| J000099072 | J000094918 | BAS | 0.010 |
| J000100452 | J000095054 | BAS | 0.001 |
| J000102119 | J000095054 | BAS | 0.001 |
| J000102503 | J000095054 | BAS | 0.001 |
| J000099196 | J000095633 | SEC | 0.001 |
| J000101155 | J000099257 | BAS | 0.001 |
| J000103716 | J000099257 | BAS | 0.001 |
| J000100809 | J000099999 | BAS | 0.001 |
| LG0830F1385P0 | TM00208 | CLA | 0.001 |
| LG1265F160P0 | TM00256 | CLA | 0.001 |
| LG1191aF2114P0 | TM00832 | CLA | 0.001 |
| LG1475F356P0189P1 | TM00939 | BAS | 0.001 |
| TM01148F198P0 | TM01148 | CLA | 0.001 |
| TM01148F200P0 | TM01148 | CLA | 0.011 |
| TM01243F578P0 | TM01243 | CLA | 0.001 |
| TM01245F588P0 | TM01245 | CLA | 0.001 |
| TM01245F589P0 | TM01245 | CLA | 0.001 |
| TM01406F1141P0 | TM01406 | PRI | 0.001 |
| TM01406F1142P0 | TM01406 | PRI | 0.001 |
| TM01448F1275P0 | TM01448 | BAS* | 0.001 |
| TM01448FPT | TM01448 | SEC* | 0.010 |
| TM01563F273P0 | TM01563 | SEC | 0.002 |
